# Heterogeneous reconstruction algorithms for cryoEM achieve limited particle classification accuracy on real benchmark datasets

**DOI:** 10.64898/2026.05.08.722747

**Authors:** Laurel F. Kinman, Andrew V. Grassetti, Maria V. Carreira, Joseph H. Davis

## Abstract

The emergence of single-particle cryoEM as a powerful method for structure determination has in large part been fueled by its ability to resolve both single static structures and complex conformational landscapes. Indeed, modern approaches to the heterogeneous reconstruction task can resolve 100s-1,000s of different maps from a single cryoEM dataset. How accurate these algorithms are, however, has proven difficult to rigorously assess, due to a lack of suitable benchmark datasets containing both realistic noise features and ground-truth labels. To address this obstacle, we recently developed a series of benchmark datasets that leverage the targeting power of Cas9 and the programmable heterogeneity of DNA to newly offer access to ground-truth per-particle structural labels in real data. Here, we challenged two popular heterogeneous reconstruction algorithms with mixed particle stacks resampled *in silico* from these datasets, finding that existing approaches resolve the encoded heterogeneity with limited accuracy. In particular, in realistic particle stacks with complex, multi-scale, and multi-axis heterogeneity, we observed that reconstruction of encoded heterogeneity depended strongly on the application of prior information about where heterogeneity was expected, and that individual particle assignments were made with significant error even when the correct structural states were reconstructed. Both molecular breathing motions and data collection features, such as defocus and projection angle, contributed to the observed particle assignment error. These results highlight important shortcomings of existing heterogeneous reconstruction methods and suggest new avenues for method development in both data collection strategies and in heterogeneous classification and reconstruction algorithms.

## INTRODUCTION

In the last decade, single-particle cryogenic electron microscopy (cryoEM) has emerged as a dominant method for determining near-atomic resolution structures of biological macromolecules (Cheng 2018). In contrast to methods like X-ray crystallography, where the recorded data is averaged across all the unit cells in a crystal, cryoEM accesses information about individual molecules (Callaway 2015). As such, cryoEM provides a uniquely powerful opportunity to resolve the complex conformational landscapes sampled by biological machines, including molecular motions that underpin environmental regulation, catalysis, and more (Amann *et al*. 2023). Heterogeneous reconstruction methods that infer multiple structures from a single dataset have thus been widely adopted in the field, and indeed the last five years have seen an explosion in the number and diversity of these approaches (Zhong *et al*. 2021; M. Chen and Ludtke 2021; Punjani and Fleet 2021, 2023; Schwab *et al*. 2024; Gilles and Singer 2025).

Early approaches to the heterogeneous reconstruction task were fundamentally discrete in nature: they assumed each particle image belonged to one of some number *k* classes and they involved the iterative sorting of particles among these classes (Scheres *et al*. 2005). Such classification-based approaches continue to be influential and widely used, particularly as implemented in cryoSPARC (Punjani *et al*. 2017), RELION (Scheres 2012), and cisTEM (Lyumkis *et al*. 2013). Nonetheless, they have substantial limitations, including that many types of structural heterogeneity are poorly described by a discrete model (Lederman and Singer 2017). Even for types of structural heterogeneity—such as compositional changes—that would seem to map well to a discrete model, classification results are often not robust to the selected value of *k*, and the true value of *k* is rarely known *a priori* (Rabuck-Gibbons *et al*. 2022).

Such limitations have led to the development of continuous models of heterogeneity. In these approaches, particles are generally mapped into a continuous latent space, from which 3D volumes can subsequently be generated (Kinman *et al*. 2023; Toader *et al*. 2023). In principle, this allows each particle to be assigned to a unique coordinate in the conformational landscape sampled by the protein, with particle images representing similar states that, ideally, co-cluster. Approaches to these particle mapping and volume generation tasks vary, including linear decomposition methods (Penczek *et al*. 2011; Tagare *et al*. 2015; Punjani and Fleet 2021; Gilles and Singer 2025) and non-linear models that leverage the power of machine learning (Zhong *et al*. 2021; M. Chen and Ludtke 2021; Punjani and Fleet 2023; Schwab *et al*. 2024). Importantly, even approaches that share a conceptual framework or architecture, like cryoDRGN (Zhong *et al*. 2021) and e2gmm (M. Chen and Ludtke 2021), can vary substantially in their implementation and exhibit differential performance.

The existence of these different approaches and implementations underscores the importance of comprehensive experimental benchmarks for algorithm performance. Such benchmarks would facilitate not only comparisons across approaches, but also the identification of each method’s failure modes, thereby enabling the iterative improvement of these methods. Previous benchmarking efforts have coalesced around a series of highly annotated real datasets, such as EMPIAR-10180 (Plaschka *et al*. 2017) and EMPIAR-10076 (Davis *et al*. 2016), where previous annotations and biochemical plausibility stand in for ground truth, as well as simulated datasets (Jeon *et al*. 2024) that offer access to ground truth but lack noise features common to real datasets. Nonetheless, there remains a substantial gap between algorithm performance on such simulated datasets and the performance routinely achieved on real datasets, which motivates the need to rigorously assess how these algorithms perform in practice (Grassetti *et al*. 2026).

Beyond enabling quantitative comparisons for method development work, obtaining realistic estimates of algorithm accuracy will be a critical first step in the path towards truly quantitative structural biology. Already, the field is moving toward more quantitative interpretations of structural heterogeneity. Beyond identifying alternative conformational states, many studies now seek to estimate their relative abundances and to compare those abundances across conditions, perturbations, or time (Haselbach *et al*. 2017; Sun *et al*. 2023; Ghanbarpour *et al*. 2023; May *et al*. 2026). Such analyses depend not only on recovering the correct set of conformational states, but also on assigning individual particle images to those states with sufficient accuracy. In practice, that assignment problem is intrinsically difficult (Evans *et al*. 2026): cryoEM particle images have low signal-to-noise ratios, multiple axes of structural variability often coexist within the same specimen, and technical parameters, such as particle pose or defocus, may influence which particles can be classified reliably. Without realistic benchmarks bearing per-particle ground truth, the magnitude and sources of these errors remain poorly defined.

To address this broader benchmarking deficiency, we recently developed a series of real datasets with encoded ground-truth heterogeneity (Grassetti *et al*. 2026). These datasets, which each contain a catalytically inactive Cas9 protein•sgRNA complex bound to a single DNA target (tDNA) of variable length, can be mixed *in silico* at prescribed frequencies and used to challenge heterogeneous reconstruction algorithms. In these datasets, not only are the true structural states—the tDNA extension lengths present in the dataset—known, but every particle image has a ground-truth “state” label (*i*.*e*., its tDNA extension length), allowing us to evaluate heterogeneous reconstruction algorithms on both how accurately they infer the correct set of structures and how accurately individual particle images are partitioned between conformational states.

Here, we used these datasets to benchmark the performance of two widely used methods, 3D classification, as implemented in cryoSPARC (Punjani *et al*. 2017) and cryoDRGN (Zhong *et al*. 2021). We found that even when the correct conformational states were reconstructed, particles were assigned to these states with limited accuracy. Confounding conformational heterogeneity, in the form of molecular breathing motions that are likely ubiquitous in real datasets, contributed to this misclassification. Interestingly, we also identified experimental imaging parameters that contributed to misclassification, highlighting the need for not only improved data processing methods but also data collection strategies tailored to the analysis of structural heterogeneity.

## RESULTS

### Large-scale heterogeneity dominates 3D classification

To better understand how existing heterogeneous reconstruction algorithms perform on real cryoEM data, we first performed 3D classification using datasets we constructed bearing encoded ground-truth heterogeneity. These datasets leverage the programmable heterogeneity of DNA, with each dataset consisting of a catalytically inactive Cas9 complexed with guide RNA and a variable-length tDNA scaffold (Grassetti *et al*. 2026). We randomly sampled an equal number of particles from each of the 13 datasets we collected, yielding a final stack with 825,864 total particles and tDNA scaffold extensions ranging in length from 2-bp to 14-bp. We subjected this stack to 3D classification in cryoSPARC using default parameters and refined each of the resulting 13 classes. We measured the occupancy of the tDNA extension in each refined volume using MAVEn (Sun *et al*. 2023) (see Methods) and found quantifiably less variation in this occupancy across the classes than when each dataset was refined individually. Furthermore, we found that particles did not partition based on their dataset of origin, with only 8.4% of particles sorted into the correct class (**Figure 1A**, see Methods). This was consistent with the notion that non-encoded heterogeneity (*i*.*e*., heterogeneity in regions outside the tDNA extension), which we observed within the consensus refinements (Grassetti *et al*. 2026), could drive per-particle classification. Indeed, inspection of the resulting class volumes highlighted large conformational changes in the REC2, REC3, and HNH domains, suggesting that 3D classification primarily sorted on non-encoded heterogeneity (**Figure 1A**).

**Figure 1.**
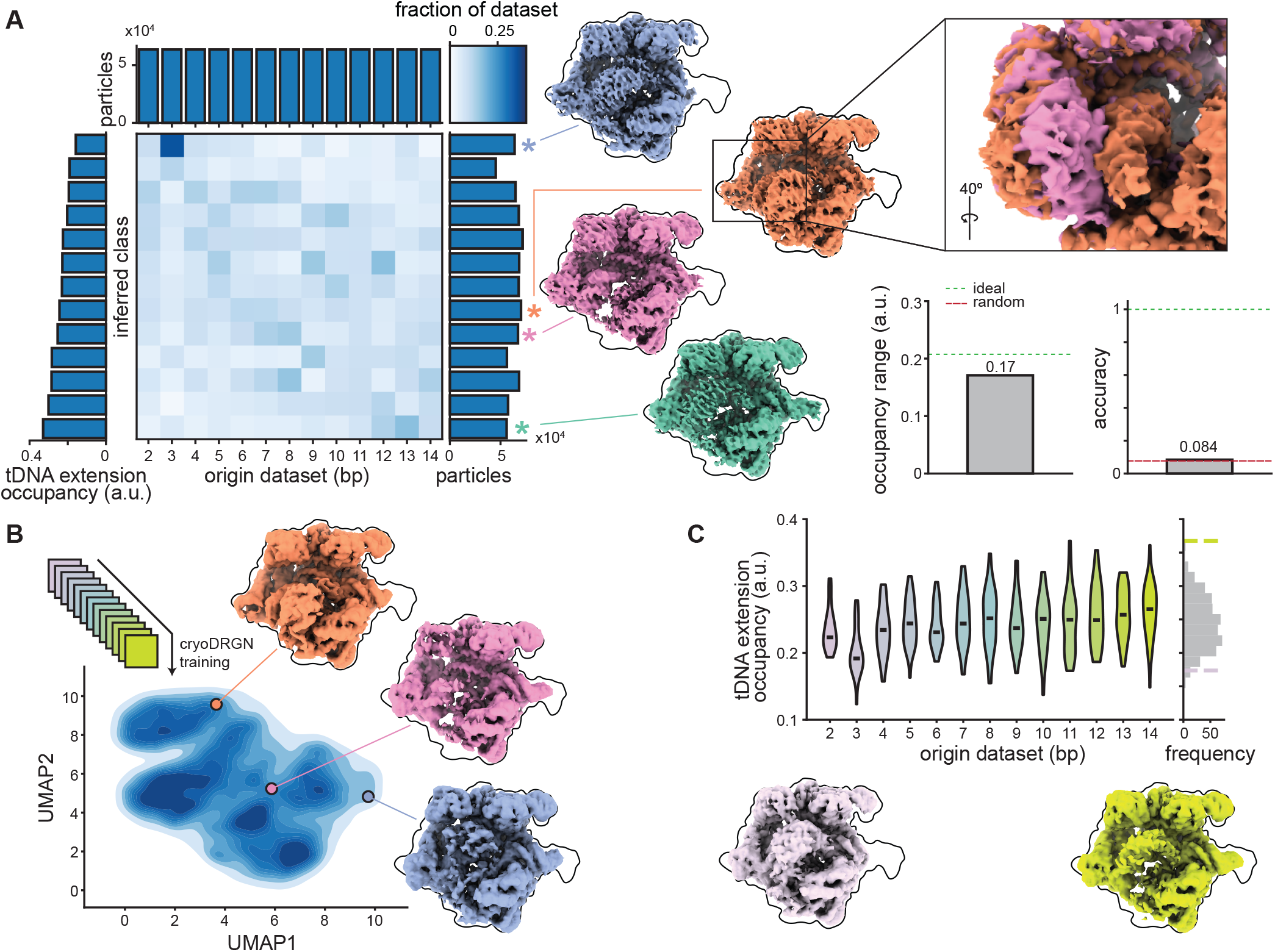
Non-encoded heterogeneity dominates outputs of heterogeneous reconstruction algorithms. **(A)** Confusion matrix of 3D classification results (without a focused mask, see Methods) on the 2-14 -bp mixed particle stack. Matrix depicts fraction of particles from each origin dataset (columns) that were assigned to inferred classes (rows), colored according to key with marginal distributions depicted. Exemplar refined maps of noted classes also depicted, including inset highlighting REC3 domain movements. Particle sorting accuracy and occupancy range of the resulting reconstructions plotted as a bar graph, with dashed lines indicating values expected from either ideal (green) or random (red) sorting. **(B)** Latent space of a cryoDRGN model trained on the 2-14 -bp mixed particle stack (see Methods), with volumes reconstructed by the cryoDRGN decoder network sampled from marked locations in latent space. **(C)** Violin plot of normalized tDNA extension occupancy measurements from 500 volumes sampled from the trained cryoDRGN model (see Methods), conditioned on origin dataset (top). Black bars depict median values for each dataset. Colored dashed lines indicate tDNA extension occupancies for representative sampled volumes, which are depicted below and colored accordingly.

Given the presence of confounding non-encoded heterogeneity within the dataset, we considered whether an approach that maps each particle image into a continuous latent space, rather than a discrete class, would better recapitulate the encoded heterogeneity. To test this, we trained a cryoDRGN model (see Methods) on our mixed particle stack and sampled volumes corresponding to individual particle images from the resulting latent space using the trained decoder network (see Methods). Much like 3D classification, these volumes primarily highlighted large-scale non-encoded heterogeneity in the REC2, REC3, and HNH domains (**Figure 1B**) overlaid with meaningful but limited variation in the length of the tDNA extension. Reflecting the limited variation observed in the tDNA extension length, volumes reconstructed from particles bearing very short (2-bp) or very long (14-bp) tDNA extensions exhibited density consistent with more intermediate extension lengths (**Figure 1C**).

### Focused 3D classification ameliorates but does not eliminate particle classification errors

We hypothesized that applying prior knowledge of the location of the encoded heterogeneity (*i*.*e*., the location of the tDNA extension) might improve classification performance. For these analyses, we used a simpler model of heterogeneity than our full stack spanning 13 datasets, instead sampling an equal number (*n*=197,780) of particles from only the datasets with a 2-bp, 8-bp, or 14-bp extension. Focused 3D classification on the tDNA extension produced three classes with the expected tDNA extension lengths: one short (2-bp), one medium (8-bp), and one long (14-bp). The difference in occupancy between the longest and the shortest tDNA extensions was similar to the occupancy range we observed when each dataset was refined individually (**Figure 2A**). Notably, this result required the focused mask. Without it, 3D classification yielded minimal variation in tDNA extension occupancy (**Supplementary Figure 1**).

**Figure 2:**
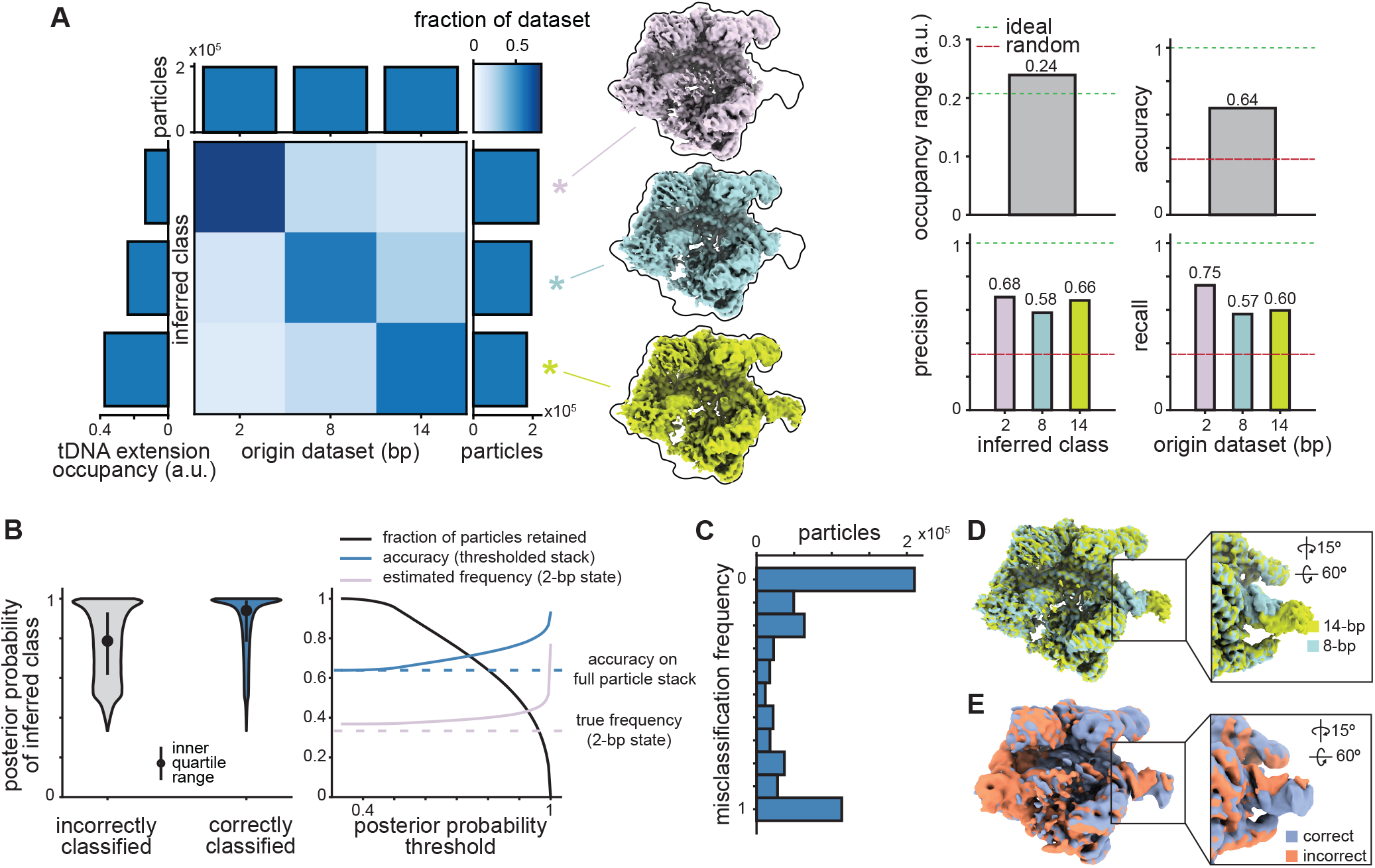
Confounding conformational and compositional heterogeneity limit particle sorting accuracy in focused 3D classification. **(A)** Confusion matrix of 3D classification results (with a focused mask, see Methods) on the 2/8/14-bp mixed particle stack. Matrix depicts fraction of particles from each origin dataset (columns) that were assigned to inferred classes (rows), colored according to key with marginal distributions depicted. Refined maps are shown for each resulting class. Occupancy range of the tDNA extension, particle sorting accuracy, and per-class precision and recall results plotted as bar graphs, with dashed lines indicating values expected from either ideal (green) or random (red) sorting. **(B)** Violin plot depicting distribution of per-particle posterior probabilities estimated by focused 3D classification, conditioned on whether particles were correctly or incorrectly classified (left). Accuracy and estimated 2-bp state frequency are plotted as a function of the lower bound imposed on the posterior probability for accepted particle assignments (right), with true frequency of the 2-bp state and accuracy on the full particle stack indicated with dashed lines. **(C)** Per-particle misclassification frequencies across 10 random seed replicates, plotted as a bar chart. **(D)** 8- and 14-bp class maps from (A), overlaid. **(E)** Refined and low-pass filtered maps from 14-bp particles that were always correctly or always incorrectly classified across ten 3D classification replicates.

Though focused 3D classification produced reconstructions largely consistent with the expected structural states, individual particle images were not classified with high accuracy. Indeed, particle sorting accuracy was only ∼64% across the entire dataset (*i*.*e*., roughly one third of the particles were errantly classified). We titrated a variety of classification parameters, including filter resolution, online expectation-maximization (O-EM) batch size, and initial learning rate during O-EM, to determine whether non-default values for these parameters would enable more accurate particle sorting (see Methods). In general, we found that use of non-default values led to equivalent or worse performance as assessed by particle sorting accuracy and occupancy range (**Supplementary Figure 2**), which we used as a surrogate for 3D density map reconstruction fidelity. However, for filter resolution specifically, which requires a user-provided value, we found that increasing this value (*i*.*e*., performing the classification at lower resolution) resulted in more accurate classification; the classifications shown here used a value of 12Å for this parameter (**Supplementary Figure 2**).

We observed several other factors that contributed significantly to accuracy. As described in Grassetti *et al*., the particle stacks used for the classifications presented here were scale-filtered and contained poses from a deep classification and refinement approach (Grassetti *et al*. 2026). Omitting either step reproducibly degraded classification accuracy (**Supplementary Figures 3-4**). Moreover, we observed a pronounced effect of particle stack size on classification accuracy, with larger particle stacks being sorted more accurately (**Supplementary Figure 5**).

Finally, to test how accurately individual particle images would be classified when correct volumes were provided, we performed a single round of supervised focused 3D classification, where particles were assigned to ground-truth volumes obtained by individually refining each dataset (see Methods). This classification achieved similar particle sorting accuracy to our previous unsupervised focused 3D classification, consistent with the idea that resolving the correct volumes is not sufficient for accurate particle classification. In contrast to unsupervised focused 3D classification, however, accuracy in this single-round assignment depended on high-resolution information (**Supplementary Figure 6**).

### Some particles are preferentially misclassified

To better understand what factors limited particle sorting accuracy, we closely examined our 3D classification results. We observed that 2-bp particles were sorted with higher accuracy (75%) than either 8-bp (57%) or 14-bp (60%) particles (**Figure 2A**), and that misclassified particles were more likely to be assigned to their class with low confidence than correctly classified particles (**Figure 2B**). This result suggested that particle sorting accuracy could be increased by only accepting class assignments above some threshold posterior probability, and indeed we observed that imposing an increasingly stringent threshold resulted in progressively higher accuracy. However, to achieve accuracy greater than 85% by this approach required a threshold that eliminated 84% of the particle stack, as many particles were misassigned with relatively high confidence. Importantly, this approach produced an artificially inflated estimate of the frequency of the 2-bp state in the particle stack due to the higher average accuracy with which 2-bp particles were classified (**Figure 2B**).

We next considered whether performing multiple replicates of the same classification could increase particle sorting accuracy, reasoning that if all particles were equally likely to be misclassified, and were more likely to be correctly classified than incorrectly classified, then averaging class assignments across multiple runs might lead to a higher effective accuracy. To test this hypothesis, we performed 10 replicates of the same focused 3D classification using different random seeds and assessed how frequently each particle was misclassified across these replicates (**Supplementary Figure 7A**). We found that particles were not equally likely to be misclassified. Instead, we observed a distinct bimodal distribution, where many particles were never misclassified but a minority of particles were always misclassified (**Figure 2C**). These misclassified particles were not simply poorly picked particles, as they produced clearly resolved 2D class averages (**Supplementary Figure 7B**).

### Conformational flexing confounds particle sorting

In comparing the refined maps from our focused 3D classification, we observed a minor change in the flexing of the tDNA extension between the 8-bp and 14-bp classes, with the 8-bp extension flexed slightly forward and the 14-bp extension flexed slightly back (**Figure 2D**). Notably, this pattern held across all replicates where an 8-bp class was successfully identified. This led us to reason that the bimodal distribution of misclassification frequency might result from confounding conformational heterogeneity within the tDNA extension. To test this, we isolated the 14-bp particles that were misclassified across all 10 replicate 3D classifications and performed a new refinement using these particles. We similarly refined the 14-bp particles that were never misclassified. In comparing the resulting maps, we found that the 14-bp particles that were always misclassified had weaker density for the tDNA extension, and that this density was flexed forward in a manner similar to the 8-bp classes (**Figure 2E-F**).

Given the confounding conformational heterogeneity introduced by bending motions of the tDNA extension we observed, we asked whether increasing the number of classes into which we sorted particles would increase the accuracy of our sorting. Briefly, we performed 3D classification using the same parameters and 2/8/14-bp mixed particle stack as before, but increased the number of classes to 9, 27, 81, or 243 total classes. As expected, this “wide classification” approach resolved more diverse flex states of the tDNA extension than in the original 3-class classification (**Figure 3A**). However, doing so did not substantially increase particle sorting accuracy: even with the largest number of classes, we did not observe the emergence of a clear trimodal distribution of tDNA extension occupancies, and even classes with the most extreme tDNA extension occupancies contained many contaminant particles (**Figure 3B**).

**Figure 3.**
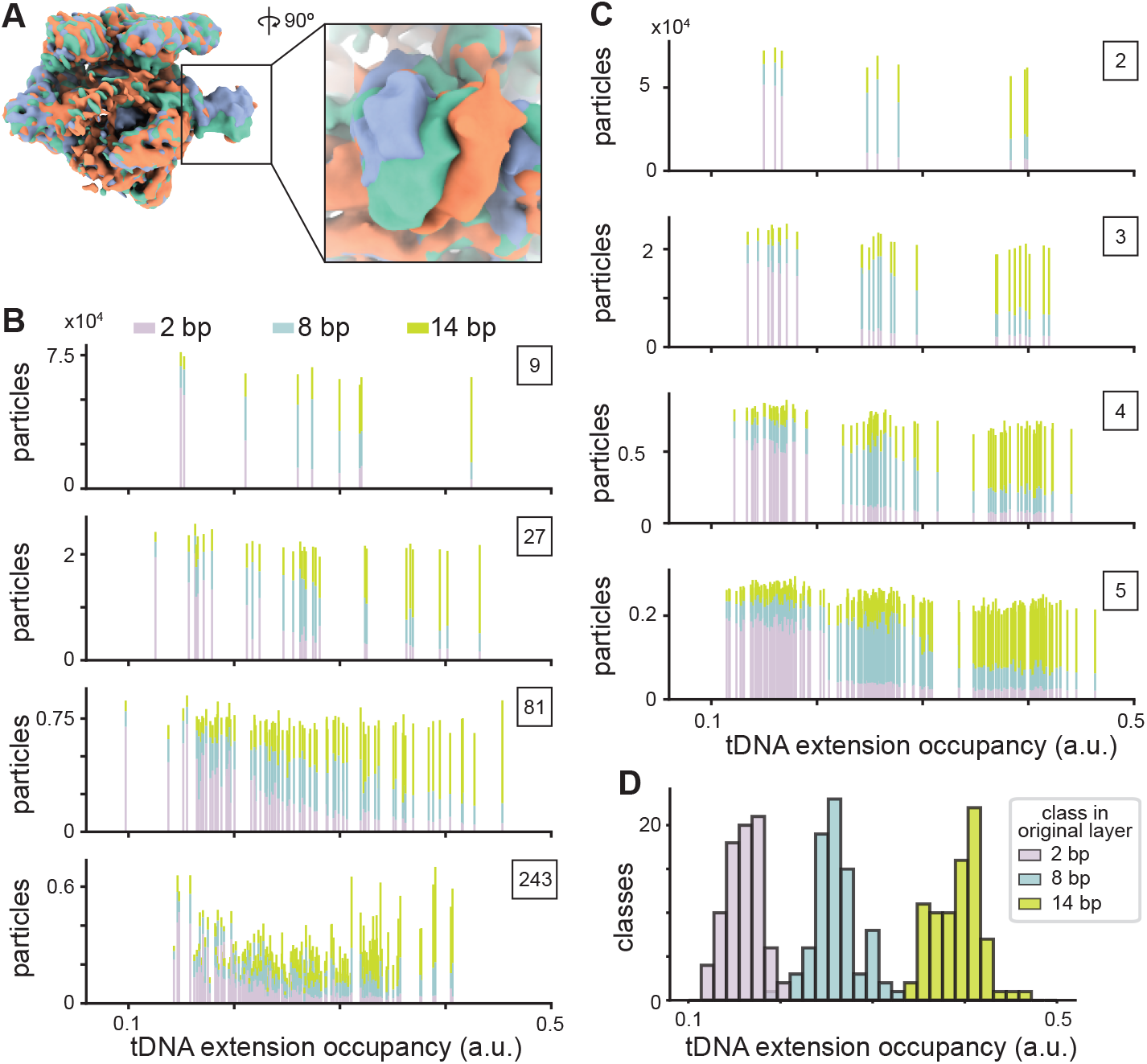
Increasing the number of classes does not markedly improve particle sorting accuracy. **(A)** Refined, low-pass filtered density maps from three representative classes exhibiting high occupancy of distinct tDNA extension states. These classes were selected to highlight a diversity of conformational flex states obtained from focused 3D classification of the 2/8/14-bp mixed particle stack using *k* = 27 total classes. **(B-C)** Stacked bar charts showing the distribution of particles assigned to each class as a function of tDNA extension occupancy. Bars are colored according to the ground-truth tDNA extension label (dataset-of-origin). Panel (B) shows results from the wide classification approach, with boxed numbers indicating the total number of classes, while (C) shows results from successive layers of deep classification, with classification layer annotated in box. Mixed-color bars highlight inaccurate particle classification. **(D)** Histogram depicting the distribution of tDNA extension occupancies in the final layer of classification for all classes that descended from the original 2-bp class (purple), the original 8-bp class (blue), or the original 14-bp class (green).

We similarly tested whether a hierarchical “deep classification” approach, where particles are progressively subclassified through multiple classification layers, would improve particle sorting accuracy. To do so, we subclassified each of the three original classes into three additional classes, and so on, producing a total of 9, 27, 81, or 243 classes at subclassification layers two, three, four, and five, respectively. As with the wide classification approach, we found that classes with very large and very small tDNA extension occupancies nonetheless retained contaminant particles (**Figure 3C**). Moreover, while this approach did produce the expected trimodal distribution of tDNA extension occupancies, this distribution was a direct result of the original classification layer. For example, particles ended up in a final class with tDNA extension occupancy greater than 0.33 if and only if they were classified as 14-bp particles in the original classification, suggesting that additional layers of classification did not successfully move 14-bp particles from the original 2-bp or 8-bp class to a final 14-bp class (**Figure 3D**). For both wide and deep classification, we found that initialization mode had little effect on classification results, as using PCA initialization instead of the default initialization produced similar results (**Supplementary Figure 8**).

### Defocus and pose impact particle sorting accuracy

In the wide and deep classification experiments, we were surprised to find 2-bp particles that were misclassified as 14-bp, as this mistake, which corresponds to a compositional change of ∼8 kDa of mass, could not be explained by a confounding conformational variable. To understand what factors were contributing to these errors, we performed focused 3D classifications as before on a particle stack consisting of only 2- and 14-bp particles. These classifications were more accurate than the classifications on the mixed stack composed of 2-, 8-, and 14-bp particles, with particle sorting accuracy between 81.5-81.6% across 10 randomly seeded replicates (**Figure 4A**). As before, misclassified particles were, on average, sorted with lower confidence than correctly classified particles (**Supplementary Figure 9**). Interestingly, we again found that particles were sorted consistently across replicates: the overwhelming majority of particles were either always correctly classified or never correctly classified (**Figure 4B**) and, again, particles that were never correctly classified were clearly resolved in 2D classes, consistent with them being true particles and not errant particle picks (**Supplementary Figure 10**).

**Figure 4.**
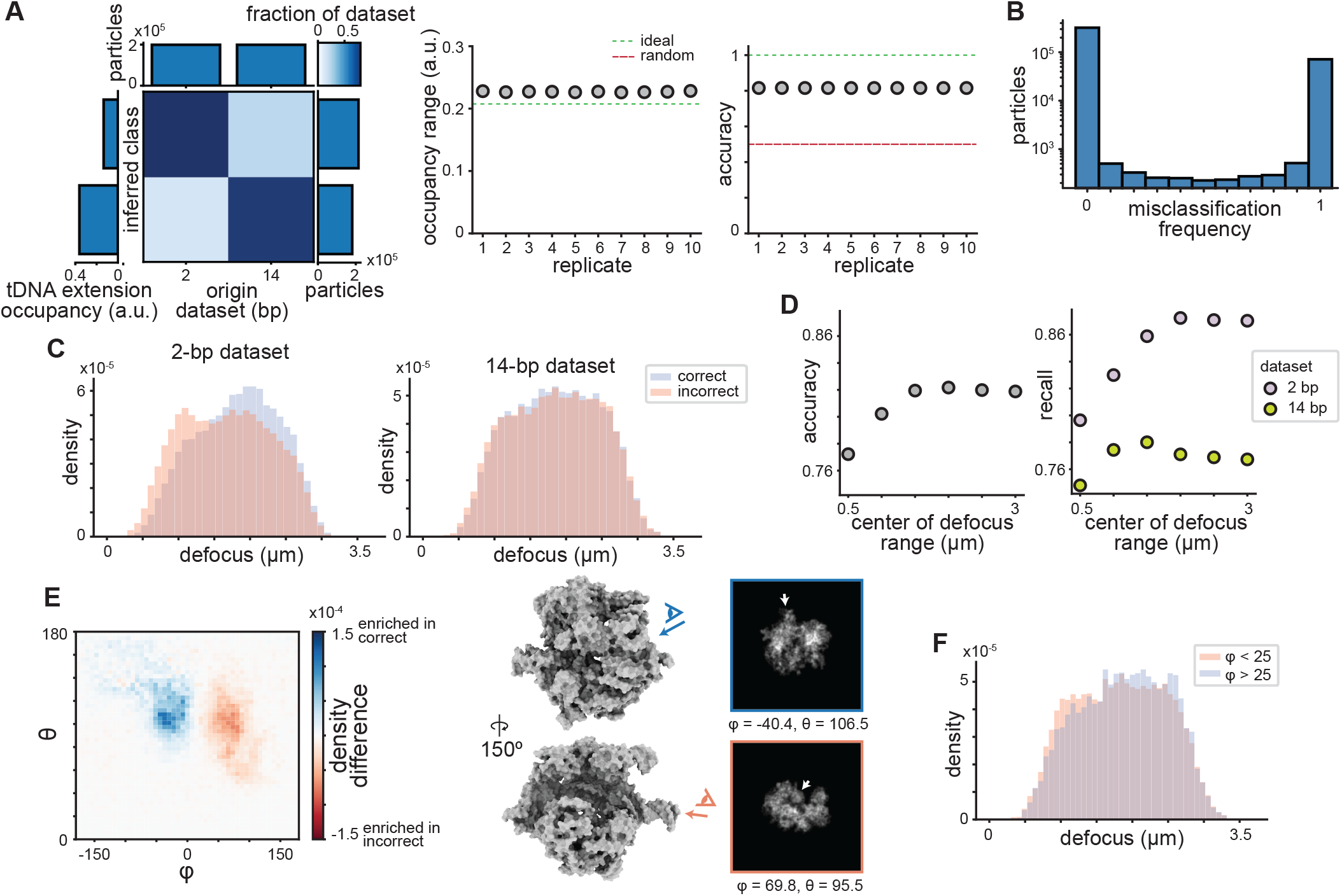
Particle assignment accuracy varies with particle pose and defocus. **(A)** Confusion matrix of 3D classification results (with a focused mask, see Methods) on the 2/14-bp mixed particle stack. For a single replicate, matrix depicts fraction of particles from each origin dataset (columns) that were assigned to inferred classes (rows), colored according to key with marginal distributions depicted. Scatter plots depict observed occupancy range and sorting accuracy across ten replicate classifications, with dashed lines indicating values expected from either ideal (green) or random (red) sorting. **(B)** Per-particle misclassification frequencies across 10 random seed replicates analyzed in (A), plotted as a bar chart. **(C)** Overlaid histograms depicting distributions of defocus values for particles in either the 2-bp (left) or 14-bp (right) dataset, conditioned on whether they were always correctly classified (blue) or never correctly classified (peach). **(D)** Scatter plot of classification accuracy (left) and recall (right) for particle stacks sampling different defocus ranges (see Methods). **(E)** Difference in viewing angle distributions for 14-bp particles that were always correctly classified or always incorrectly classified (left). The viewing angles most enriched in correct and incorrect particles, respectively, are shown alongside the atomic model and corresponding projection images. White arrowheads indicate the position of the tDNA extension. **(F)** Overlaid histograms depicting distribution of defocus values of incorrectly classified 14-bp particles, conditioned on value of the pose angle φ.

This observation suggested that some particle images are hard to classify even without confounding conformational heterogeneity. To better understand what image features could make a particle hard to classify, we compared metadata from particles that were always correctly classified to metadata from particles that were never correctly classified. In doing so, we found that the misclassified 2-bp particles were enriched for images collected close to focus. In contrast, there was no apparent preference for close-to-focus particles in the misclassified 14-bp particles (**Figure 4C**). Comparing the final per-particle defocus estimates obtained after CTF refinement and Bayesian polishing to the original micrograph-level defocus estimates indicated that per-particle defocus estimates largely agreed with micrograph-level defocus estimates, with no clear correlation between the degree of defocus estimate divergence and whether the particle was correctly or incorrectly classified (**Supplementary Figure 11**). These results suggested that particles collected at high defocus might be easier to classify, consistent with the higher contrast and increased low-resolution signal present in these images (Sigworth 2016). To test this hypothesis, we subsampled this particle stack further, creating smaller particle stacks composed of equal numbers of 2-bp and 14-bp particles within a defined defocus range (see Methods). We performed focused 3D classification on each of these stacks and assessed the resulting particle sorting accuracy, finding that accuracy increased with image defocus, and that this increase was driven primarily – although not exclusively – by more accurate sorting of the 2-bp particles (**Figure 4D**).

While we did not observe a strong relationship between defocus and particle sorting accuracy in the 14-bp dataset, we did uncover a difference in the pose distribution of correctly and incorrectly classified 14-bp particles. Specifically, the misclassified 14-bp particles were highly enriched for a population of projection angles (φ > 25) observed at only very low frequency in the correctly classified particles. These projection angles correspond to a view roughly parallel to the tDNA extension (**Figure 4E**). 2D classification of misclassified particles from populations with different viewing angles suggested that they represented two *bona fide* viewing angles, rather than a minority population of particles with poor projection angle estimates (**Supplementary Figure 12**). Finally, when we visualized the defocus distribution again, this time conditioned on the viewing angle, we found that misclassified 14-bp particles with the tDNA extension orthogonal to the projection angle (φ < 25) exhibited an enrichment for low defocus values (**Figure 4F**), consistent with both pose and defocus contributing to misclassification in the 14-bp dataset.

### Continuous models of heterogeneity do not achieve high accuracy particle sorting

Having observed that 3D classification achieved only moderate particle sorting accuracy on the 2/8/14-bp mixed particle stack, and that this accuracy was limited by factors including confounding conformational heterogeneity, we sought to assess whether a continuous model like cryoDRGN would sort particles more accurately. To test this, we trained a cryoDRGN model using the 2/8/14-bp mixed particle stack and subsequently sampled a 3D volume corresponding to each particle (Sun *et al*. 2023). We measured the occupancy of the tDNA extension in each of these volumes using MAVEn (see Methods) and found both that cryoDRGN successfully reconstructed volumes with diverse tDNA extension lengths and that tDNA extension length in the reconstructed volume correlated with the origin dataset of the corresponding particle image (**Figure 5A**). We then assigned particles to three classes (2-bp, 8-bp, or 14-bp) based on their tDNA extension lengths (see Methods) and found that cryoDRGN accurately assigned ∼52% of particles to their correct conformational state (**Supplementary Figure 13**). While this accuracy was much higher than that observed in a comparable discrete classification (*i*.*e*., without a focused mask – 38%), it was lower than the accuracy achieved through focused 3D classification (64%). To test whether applying a mask would increase particle sorting accuracy in cryoDRGN, we generated a masked particle stack (see Methods) that we used to train a new cryoDRGN model (**Figure 5B**). We assessed the accuracy of this cryoDRGN model as before, and found that, unlike in 3D classification, applying a mask in cryoDRGN only modestly improved particle sorting accuracy (to 58%) (**Figure 5C, Supplementary Figure 14**). This effect was not unique to cryoDRGN as supplying a focused mask on the tDNA extension also only modestly benefited classification accuracy by 3D Variability Analysis (3DVA), another continuous classification and 3D reconstruction approach (Punjani and Fleet 2021) (**Supplementary Figure 15**).

**Figure 5.**
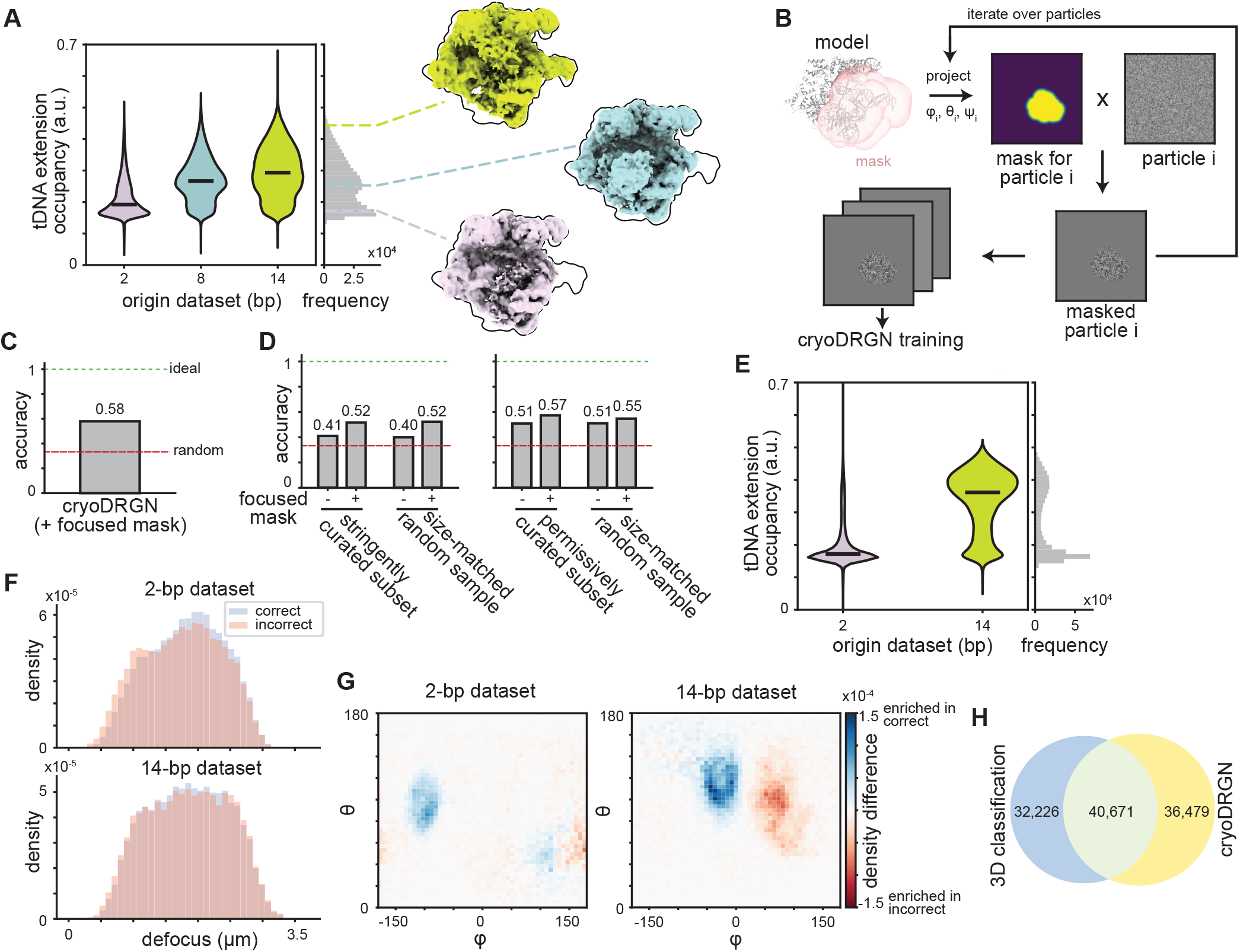
cryoDRGN outperforms unfocused 3D classification but is also challenged by close-to-focus imaging and specific poses. **(A)** Violin plot of tDNA extension occupancies recorded for every particle in the 2/8/14-bp mixed particle stack following reconstruction with the trained cryoDRGN decoder network (see Methods). Colored dashed lines indicate tDNA extension occupancies of sampled representative volumes (middle). **(B)** Schematic of the approach to mask particles for focused analysis in cryoDRGN (see Methods). **(C)** Accuracy of the cryoDRGN model trained on the masked 2/8/14-bp mixed particle stack, with ideal (green) and random (red) performance noted by dotted lines. **(D)** Accuracy of cryoDRGN models trained on the permissively or stringently curated subsets of the 2/8/14-bp mixed particle stack, with or without masking. Random and ideal performance noted as in (C). **(E)** Normalized tDNA extension occupancies recorded for each particle in the 2/14-bp mixed particle stack following reconstruction with the trained cryoDRGN decoder network. **(F)** Defocus distributions for particles in each dataset that were correctly or incorrectly classified by cryoDRGN. **(G)** Difference in viewing angle distributions between particles in each dataset that were correctly or incorrectly classified by cryoDRGN. **(H)** Venn diagram comparing particles from the 2/14-bp mixed particle stack that were always incorrectly classified by focused 3D classification or cryoDRGN.

To understand whether the presence of non-encoded heterogeneity affected cryoDRGN’s ability to accurately sort particles, we turned to the curated particle stacks described previously (Grassetti *et al*. 2026). Briefly, the full particle stack, containing all 2-14 -bp particles, was subjected to deep classification; following the final layer of classification, selected classes were pooled and refined. This approach was used to generate two particle stacks depleted of non-encoded heterogeneity to variable extent: one “permissively curated” particle stack in which a single axis of variation (motions of the REC3 domain) was depleted (*n* = 231,960 total particles), and one “stringently curated” particle stack in which all three primary axes of variation (motions of the REC2, REC3, and HNH domains) were depleted (*n* = 46,932 total particles). We trained a cryoDRGN model on each of these particle stacks, with or without masking, and assessed particle sorting accuracy in each trained model. As we previously found that particle stack size impacted particle sorting accuracy in 3D classification (**Supplementary Figure 5**), we also generated “size-matched” controls, where an equal number of particles was randomly subsampled from the 2/8/14-bp mixed particle stack. The distribution of 2-, 8-, and 14-bp particles was fixed at 1:1:1 across all stacks.

Consistent with our results from focused 3D classification, we found that particle stack size impacted particle sorting accuracy, with the smaller size-matched controls performing worse than the full stack (**Figure 5D, Supplementary Figure 14**). Interestingly, we observed no effect of depleting non-encoded heterogeneity in either dataset, suggesting that limited particle sorting accuracy does not reflect some limited representation capacity of the network (**Figure 5D, Supplementary Figure 14**). This result stands in contrast to 3D classification, where we found that depleting non-encoded heterogeneity did provide some benefit, particularly in the absence of a focused mask (**Supplementary Figure 16**). However, even in 3D classification, this benefit was present only in the curated stack and was not sufficient to offset the decrease in performance resulting from a smaller particle stack.

### cryoDRGN and 3D classification misclassify the same particles

In all cryoDRGN models we trained, we found that the distribution of the tDNA extension occupancies was generally bimodal, with 8- and 14-bp particles comingled in the high-occupancy class (**Figure 5A, Supplementary Figure 13**). In support of the difficulty in distinguishing the 8- and 14-bp particles, we found that a cryoDRGN model trained on only 197,780 2-bp and 197,780 14-bp particles with no masking achieved comparable accuracy to discrete focused 3D classification performed on that same particle stack (80.5%, **Figure 5E**). As with focused 3D classification, we sought to understand what factors limited particle sorting accuracy in this stack by comparing images that were correctly classified to those that were incorrectly classified. Our findings were largely consistent with what we observed in focused 3D classification: pose contributed strongly to misclassification, with inaccurately classified 14-bp particles again strongly enriched for the tDNA extension parallel viewing angle and depleted for the tDNA orthogonal viewing angle (**Figure 5G**). Interestingly, we also observed an effect of pose in the 2-bp dataset, with the two predominant viewing angles highly depleted in the inaccurately classified particles (**Figure 5G**). In comparison, defocus contributed weakly to misclassification in the trained cryoDRGN model, and an effect of defocus was only discernible in the 2-bp dataset (**Figure 5F**). These results suggested that cryoDRGN and 3D classification were challenged by the same subset of particles and indeed we find that there is substantial overlap between the particles that are incorrectly classified by the two approaches (**Figure 5H**).

## DISCUSSION

Motivated by the need to assess the performance of heterogeneous reconstruction algorithms on datasets with realistic noise features, we recently developed a series of experimental cryoEM datasets bearing encoded heterogeneity (Grassetti *et al*. 2026). Here, we demonstrated the power of using these datasets to benchmark algorithm performance for two widely used heterogeneous reconstruction methods, examining both how accurately these methods inferred the correct set of conformational states from the data and how accurately they mapped individual particle images to their corresponding conformational state. This benchmarking revealed important limitations of these methods, including their dependence on prior information (*i*.*e*., a mask focused on the region of interest) and the confounding effects of conformational and compositional heterogeneity. They also suggested that new software development may be insufficient to address some of these limitations, raising questions about how to better collect data to facilitate accurate heterogeneous reconstruction.

### Performant classification approaches rely on prior information

One key factor in successfully resolving the encoded heterogeneity in these datasets was the application of prior knowledge about where in the volume heterogeneity was expected: naïve 3D classification without a focused mask on the tDNA extension substantially dampened the observed variation in the extension and resulted in near-random sorting of particles relative to their dataset of origin (**Figure 1, Supplementary Figure 1**). In comparison, focused 3D classification on the tDNA extension resulted in the most accurate particle sorting across all methods tested (**Figure 2A, Figure 4A**). This requirement for prior knowledge of where heterogeneity is expected is a significant limitation of 3D classification: ideally, heterogeneity should be identifiable and quantifiable even when its location is not known in advance. Interestingly, neither of the continuous models of heterogeneity we tested displayed this requirement. Both cryoDRGN and 3DVA substantially outperformed naïve 3D classification at inferring a distribution of tDNA extension lengths and sorting particles accordingly. Taken together, these results suggest that continuous models of heterogeneity may be particularly useful for identifying regions of variability and building masks that can then be used in focused 3D classification for more accurate particle assignment (Kinman *et al*. 2025).

### Molecular breathing motions confound heterogeneous reconstruction algorithms

In both focused 3D classification and cryoDRGN, we found that even when the correct conformational states were recovered, particles were assigned to those states with limited accuracy. The inaccuracy we observed in per-particle assignments is consistent with previous theoretical work arguing that per-particle assignment is likely to be challenging due to the low SNR of the particle images and the need to estimate both pose and 3D class (Evans *et al*. 2026; Dingeldein *et al*. 2025). Nonetheless, error in per-particle assignments represents a significant hurdle for efforts to use cryoEM to infer quantitative conformational landscapes (Frank 2018).

In our datasets, one important contributor to this error was flexing of the tDNA extension. Across both methods, 2-bp particles were sorted more accurately than 8-bp or 14-bp particles, and cryoDRGN in particular struggled to distinguish between 8-bp and 14-bp particles (**Figure 2A, Figure 5A**). Meanwhile, focused 3D classification appeared to prioritize the flex state of the DNA extension in sorting particle images (**Figure 2E**); providing more classes, which we hypothesized could account for confounding conformational heterogeneity, did not boost particle sorting accuracy (**Figure 3B-C**). Together, these results suggest that multi-axis heterogeneity of this kind is difficult for existing methods to disentangle. This limitation is important because molecular breathing motions are likely widespread in real cryoEM datasets and may confound particle sorting more than previously appreciated, motivating the development of new methods that can better model such compound structural changes.

### Features of data collection make some particles difficult to classify

Another significant factor limiting particle sorting accuracy in our experiments was features of data collection itself. In examining particles that were repeatedly misassigned by focused 3D classification, we found that these particles were enriched for images collected close to focus (**Figure 4C-D,F**) and, in the 14-bp dataset, at viewing angles where the tDNA extension projected along the z-axis (**Figure 4E**). These same features also contributed to misclassification by cryoDRGN, with substantial overlap between the particles misclassified by the two approaches (**Figure 5F-H**). This latter observation is interesting because it suggests low image contrast and particular particle orientations may pose general challenges for heterogeneous reconstruction algorithms. It also forecloses on the possibility of using cryoDRGN as an orthogonal approach to correct errors made by focused 3D classification, or vice versa. We also found that particle stack size had a significant impact on the accuracy of both methods, with performance dropping precipitously on stacks with fewer than 100,000 particles (**Figure 5D, Supplementary Figure 5**). Together, these results highlight the important roles that data collection strategies play in accurate analysis of heterogeneous cryoEM datasets and motivate tailoring data collection to specifically facilitate heterogeneity analysis. Strategies for doing so might include: applying larger defocus or a phase plate to boost signal in the low spatial frequency regime (Danev *et al*. 2014; Danev and Baumeister 2016; Schwartz *et al*. 2019; Remis *et al*. 2024; Axelrod *et al*. 2024); revisiting once common orthogonal tilt methods (Rosenthal and Henderson 2003; Leschziner 2010) to obtain multiple related views of a given particle; and collecting larger datasets, even when additional particles yield marginal improvements in the resolution of the consensus refinement.

### Estimating conformational state frequencies with cryoEM

Despite significant (∼20-40%) error in per-particle assignments, the conformational state frequencies estimated in our experiments were often close to the ground-truth frequencies in the mixed particle stacks (**Figure 2A**). Indeed, in some cases, we found that optimizing for accurate per-particle assignment was counter-productive, causing the conformational state frequency estimate to deviate from the uniform state sampling tested here (**Figure 2B**). Although initially surprising, this result is consistent with recent work suggesting that using cryoEM data to refine an estimate on the distribution of states, rather than assigning and counting individual particle images, may prove to be a more accurate approach for quantifying conformational landscapes (Tang *et al*. 2023). The experimental datasets we developed and deployed for benchmarking here represent a powerful new tool for rigorously testing this hypothesis, as they can be readily resampled *in silico* at defined ratios and the ground-truth per-particle conformational state labels facilitate straightforward comparison of the quantitative accuracy of particle-counting and ensemble reweighting approaches.

## DATA AVAILABILITY

Raw movies and particle stacks used for benchmarking will be deposited at EMPIAR upon or before publication. Results from 3D classification jobs and 3DVA will be available at Zenodo upon or before publication. Weights and latent embeddings from trained cryoDRGN models are available by request.

## CODE AVAILABILITY

The cryoDRGN version used for model training is available at github.com/mariacarreira/cryodrgn_noisy_encoder. Scripts and mask used for generating the masked particle image stacks are available at github.com/lkinman/mask_particle_images. Scripts used for subsampling particle images are available at github.com/lkinman/benchmark_datasets. CryoSRPNT is available at github.com/bpowell122/cryoSRPNT.

## ACKNOWLEDGEMENTS

This research was supported by the NIH grant R01GM144542, NSF CAREER grant 2046778, and the Smith Family Odyssey Award (JHD). Molecular graphics and analyses performed with UCSF ChimeraX, developed by the Resource for Biocomputing, Visualization, and Informatics at the University of California, San Francisco, with support from National Institutes of Health R01GM129325 and the Office of Cyber Infrastructure and Computational Biology, NIAID. We thank H. Bridges, A. Punjani, and S. Ludtke for helpful discussions.

## AUTHOR CONTRIBUTIONS

Conceptualization: LFK, AVG, JHD.

Data curation: LFK, AVG.

Investigation: LFK.

Software: LFK, MVC.

Supervision: JHD.

Visualization: LFK, JHD.

Writing – original draft: LFK.

Writing – review & editing: All authors.

Funding acquisition and project administration: JHD.

## MATERIALS AND METHODS

### Generation of subsampled and curated particle stacks

Sample preparation, including application to graphene-coated grids (Grassetti *et al*., 2023), imaging, and data processing for the dCas9 datasets are described in Grassetti *et al*., 2026. For all experiments requiring random subsets, these subsets were generated using the `generate_ distributions.py` script described in the same publication. All random subsets described here contained equal numbers of particles at every included extension length (2-14 -bp, 2/8/14-bp, or 2/14-bp). For the experiments using curated subsets depleted of non-encoded heterogeneity, particles were subsampled at equal frequency for each included tDNA extension length from the curated stacks described in Grassetti *et al*. Size-matched controls were generated by subsampling the same number of particles randomly from the uncurated stack using the `generate_distributions.py` script, as described above. Final particle stack sizes were 46,932 total particles for the stringently curated subset and size-matched control, and 231,960 total particles for the permissively curated subset and size-matched control.

For the defocus titration experiments, a series of overlapping defocus range windows was defined. Each window spanned 1 μm defocus total; windows were centered on the following defocus values: 0.5, 1.0, 1.5, 2.0, 2.5, and 3.0 μm. The set of all particles in the 2/14-bp mixed stack with defocus within a given window was subsampled to produce a final stack with 19,000 2-bp particles and 19,000 14-bp particles whose defocus was in the defined range. These subsampled stacks were subjected to 3D classification and assessment of classification accuracy as described below. Finally, for the particle stack size titration experiments, the desired number of particles were subsampled from the 2/8/14-bp mixed stack using the `generate_distributions.py` script. Each subsample contained a 1:1:1 mixture of 2, 8, and 14-bp particles. This sampling procedure was repeated to produce 10 independently sampled replicates at each stack size tested. The total stack sizes tested were: 6,000; 15,000; 30,000; 45,000; 60,000; 90,000; 120,000; 150,000; 180,000; 240,000; 270,000; 300,000; 360,000; 420,000; 480,000; 540,000; and 600,000 particles. Each stack was independently imported into cryoSPARC and classified as described below.

### 3D classification in cryoSPARC

3D classification was performed in cryoSPARC without pose refinement. Poses used in 3D classification resulted from the deep classification and pooled refinement approach described in Grassetti *et al*., 2026. All 3D classification jobs used a solvent mask generated as follows. Using the atomic model of the 14-bp tDNA·sgRNA·dCas9 complex (Grassetti *et al*., 2026), we simulated a density map at 8Å resolution using the molmap tool in ChimeraX (Meng *et al*. 2023). This map was summed with the RELION-refined map of the 2/8/14-bp mixed particle stack (Grassetti *et al*., 2026), and the composite map was resampled on the grid of the refined map in ChimeraX. The resulting map was imported into cryoSPARC, where the following modulations were applied: a low-pass filter (15Å), a dilation of 12 pixels, a soft-padding width of 12 pixels, and binarization. For focused 3D classification, a focused mask on the tDNA extension was generated similarly: the ChimeraX molmap tool was used to generate a map from the final 17-bp of the 14-bp tDNA extension, which was resampled on the grid of the refined 2/8/14-bp mixed stack map. This map was low-pass filtered, dilated, soft-padded, and binarized in cryoSPARC using the same parameters as the solvent mask. Unless otherwise indicated, all 3D classification jobs used default cryoSPARC parameters and 12Å filter resolution. For parameter sweeps, all parameters except the one being tested were held at default values. For O-EM batch size and initial learning rate sweeps, the filter resolution was set at 12Å.

Wide and deep classification experiments were similarly carried out with default parameters and 12Å filter resolution. For the wide classification experiments, the only adjusted parameter was the number of classes (*k* = 9, 27, 81, and 243); for the deep classification experiments, each class from the original 3-class classification was subsequently subclassified into 3 more classes until a total of 9, 27, 81, or 243 classes was achieved. Wide and deep classifications were performed with either simple initialization or PCA initialization. For the wide classification experiments with PCA initialization, 300 reconstructions were used for PCA. To compare supervised and unsupervised classification, the same classification parameters were used as before (“unsupervised”) but the refined map from each individual dataset (2-bp, 8-bp, and 14-bp, or 2-bp and 14-bp) was provided as an input volume and particles were subjected to 0 O-EM epochs and only 1 full EM iteration. These supervised classifications were performed at both 12Å and 1Å filter resolution.

### Evaluation of 3D classification results

Following 3D classification, each class volume was independently refined, using the class volume as the initial model. The occupancy of the tDNA extension and the constant C-terminal domain (J. S. Chen *et al*. 2017) was measured as follows: first, the molmap tool in ChimeraX was used to generate maps corresponding to the last 17-bp of the tDNA extension and to the C-terminal domain (residues 1099-1368) at 3Å resolution. These maps were resampled on the grid of the refined class volume maps and binarized directly in ChimeraX to produce masks for measuring occupancy. Each refined volume from a given classification job was then queried for occupancy of the tDNA extension and C-terminal domain using MAVEn (Sun *et al*. 2023); the classes were then ranked by their normalized tDNA extension occupancy (*i*.*e*., occupancy of the tDNA extension relative to the C-terminal domain) to identify classes corresponding to each origin dataset. For example, in a 3-class classification on the 2/8/14-bp mixed stack, the class with the lowest tDNA extension occupancy would be designated the 2-bp class, the next lowest the 8-bp class, and the highest the 14-bp class. For each particle, its origin dataset was then compared to this inferred class assignment. Accuracy was calculated as the fraction of particles assigned to the correct class. Precision was calculated as the fraction of particles in an inferred class that were correctly assigned to that class, whereas recall was calculated as the fraction of particles from an origin dataset that were correctly assigned to the corresponding class. Occupancy range reflected the difference in normalized occupancy between the highest occupancy class and the lowest occupancy class. Reference values for occupancy range were obtained by similarly calculating the difference in the normalized tDNA extension occupancy for the refined map from each of the 2-bp and 14-bp datasets. For the wide and deep classifications specifically, class volumes were not refined prior to measuring occupancy. Instead, masks were generated similarly in ChimeraX, using molmap at 11Å resolution and resampling on the grid of the class volume (box size = 56 pixels), and class volumes were directly queried for occupancy of the tDNA extension and C-terminal domain.

### cryoDRGN training

cryoDRGN models were trained using a new cryoDRGN architecture (github.com/mariacarreira/cryodrgn_noisy_encoder) adapted from cryoDRGN v3.4, in which Gaussian noise, sampled from *N*(0, 1), was added to each particle’s latent embeddings during the first five epochs of training. In practice, we have found these noise injections ameliorate minor artifacts we have observed in the reconstructed density maps, and improve learned heterogeneity. All models were trained at a box size of 128 pixels (2.24Å/pix) with 8 latent dimensions, a batch size of 256, and encoder and decoder dimensionality of 1024. The same poses from deep classification and pooled refinement used in 3D classification were used for cryoDRGN training (see above). All models except those trained on the curated subset and its size-matched control were trained for 50 epochs; given the small number of particles in these stacks (46,932 particles in each), models trained on them were trained for 100 epochs.

### Generation of masked particle stacks for cryoDRGN training

Masking in cryoDRGN was executed by generating a stack of masked particles, and then training a cryoDRGN model as described above on that stack of masked particles. The mask used in cryoDRGN training covered both the tDNA extension and the dCas9 C-terminal domain and was initially generated in ChimeraX using the molmap command at 8Å resolution, then resampled on the grid of the original refined RELION map from the 2/8/14-bp mixed particle stack. In cryoSPARC, this map was resampled to a box size of 128 pixels to match the box size used in cryoDRGN training, and low-pass filtered to 15Å. The map was binarized, and dilation and soft padding were applied (radius/width 6 pixels). The `project3d.py` script implemented in cryoSRPNT (github.com/bpowell122/cryoSRPNT) was then used to generate projections of this mask matching the projection angles of the particle images in each stack (Powell *et al*., 2024). The projected mask and particle images were then multiplied to produce masked particle stacks suitable for training cryoDRGN.

### cryoDRGN model evaluation

Performance of the cryoDRGN model was assessed by measuring tDNA extension occupancy of volumes reconstructed by the cryoDRGN decoder network. For the full mixed stack, with tDNA extension lengths ranging from 2-bp to 14-bp, *k*-means clustering on the latent embeddings was used to sample and reconstruct 500 volumes at box size 128 (2.24Å/pixel). Volumes were binarized using the threshold predicted by SIREn (Kinman *et al*. 2025) and normalized tDNA occupancy was measured using MAVEn, with masks generated as described above using the molmap tool in ChimeraX, at a resolution of 4Å, and subsequently binarized. For each map, tDNA extension occupancy was normalized by the occupancy of the constant C-terminal domain. For all other cryoDRGN models, occupancy was measured “on-the-fly” at box size 64, as described previously (Sun *et al*. 2023). Briefly, masks for the tDNA extension and the C-terminal domain were generated in ChimeraX using the molmap tool at 9Å resolution and then binarized. For every particle in the stack, the corresponding volume was reconstructed by cryoDRGN, binarized at the threshold predicted by SIREn, and the occupancy of the binarized map’s tDNA extension and C-terminal domain was measured. All tDNA extension occupancy measurements were normalized by occupancy of the constant C-terminal domain. To quantify particle sorting accuracy, particles were sorted into three equal-sized classes by rank-ordering of the normalized tDNA extension occupancies. Accuracy was then calculated as the fraction of particles assigned to the correct class based on their dataset of origin. To determine robustness of accuracy calculations to the occupancy thresholds used to separate particles into classes, we calculated the effective accuracy across a wide array of threshold combinations, simultaneously titrating the lower threshold separating the 2-bp class from the 8-bp class and the upper threshold separating the 8-bp class from the 14-bp class. At each unique threshold combination, we calculated accuracy as before by determining the fraction of particles for which the inferred class matched the origin dataset.

### Analysis of misclassified particles

Comparison of correctly and incorrectly classified particles was accomplished using random-seed replicates of focused 3D classification for both the 2/8/14-bp mixed particle stack and the 2/14-bp mixed particle stack. The misclassification frequency across 10 replicates was calculated for each particle, and subsets of particles that were always (misclassification frequency = 1) incorrectly classified or never (misclassification frequency

= 0) incorrectly classified were identified. The correctly and incorrectly classified particles were further segregated by dataset of origin and reimported into cryoSPARC for 2D classification and homogeneous refinement. Metadata comparisons were performed by extracting defocus and pose data for each particle from relevant .cs and .star files. Poses were converted where appropriate to Euler angles following established conventions. Pose enrichment heatmaps were calculated by creating 2D probability histograms of the φ/θ Euler angles (disregarding the in-plane rotation, described by ψ) in each of the incorrectly and correctly classified particle populations. The represented heatmaps depict the difference of these two probability histograms. Analogous analyses were performed for particles correctly and incorrectly classified by the trained cryoDRGN model.

### 3D variability analysis

3D Variability Analysis (3DVA; Punjani and Fleet 2021) was performed on the 2/8/14-bp mixed particle stack. For the “unmasked” 3DVA experiments, the same solvent mask used for all 3D classification jobs was supplied. For the “masked” 3DVA experiments, the same focused mask over the tDNA extension that was used in focused 3D classification was supplied. In all cases, 3DVA was performed using 3 components, 12Å filter resolution, and default parameters. To assess the accuracy of 3DVA, a volume was generated “on-the-fly” for each particle in the stack, using the mean map, component maps, and the particle’s component values. The occupancy of the tDNA extension and the dCas9 C-terminal domain was measured in each of the resulting maps, using MAVEn and the same masks employed to measure occupancy of refined class volumes following 3D classification in cryoSPARC. As in cryoDRGN, particle sorting accuracy was quantified by sorting particles into three equal-sized classes based on rank-ordering of normalized tDNA extension occupancies and subsequently calculating the fraction of particles whose inferred class matched their origin dataset.

## Supplementary Information

### SUPPLEMENTARY FIGURES

**Supplementary Figure 1.**
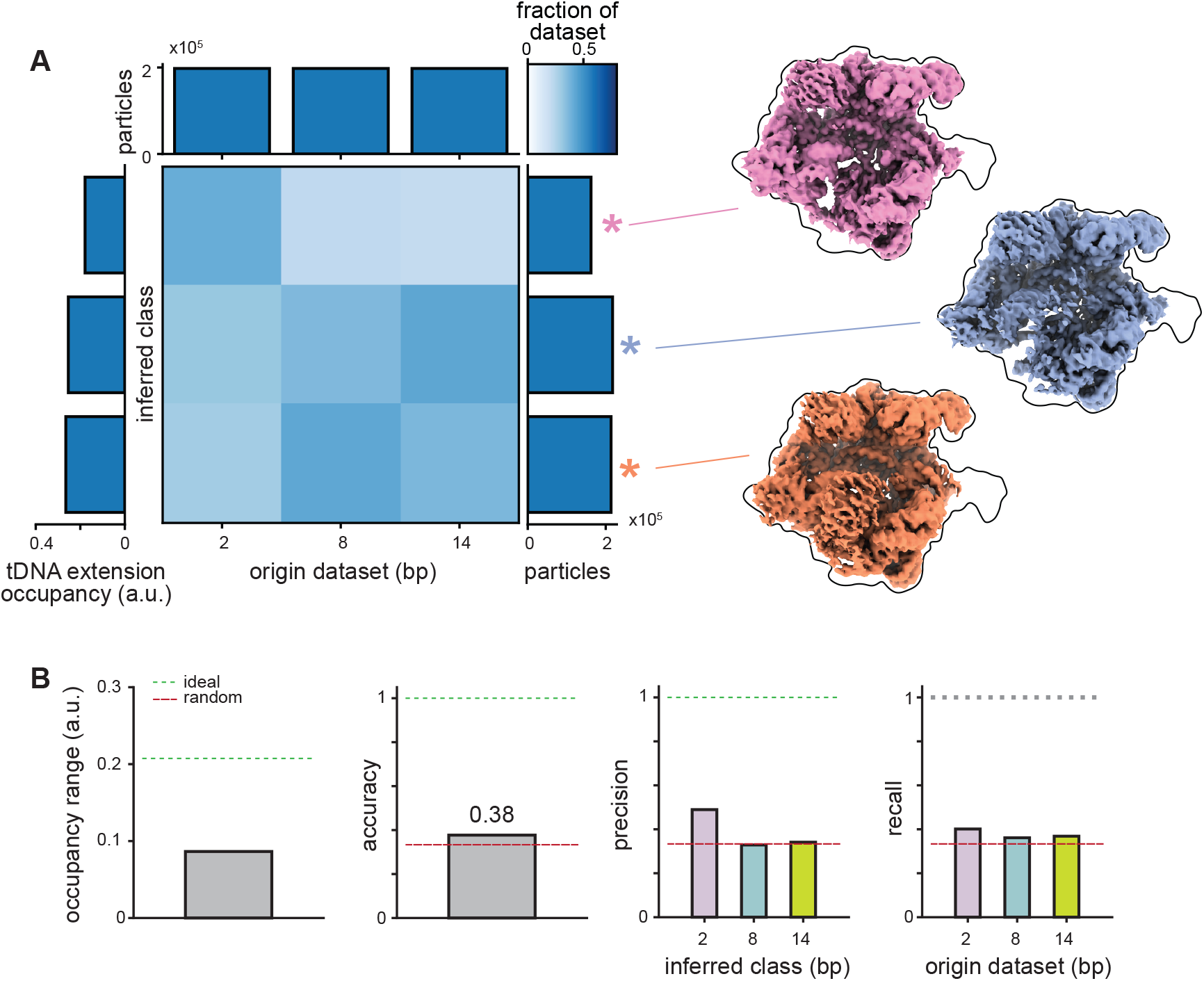
Classification performance on the 2/8/14-bp mixed particle stack without a focused mask. **(A)** Confusion matrix of global (*i*.*e*., unmasked) 3D classification results on the 2/8/14-bp mixed particle stack. Matrix depicts fraction of particles from each origin dataset (columns) that were assigned to inferred classes (rows), colored according to key with marginal distributions depicted. Refined maps are shown for each resulting class. **(B)** Occupancy range of the tDNA extension, particle sorting accuracy, and per-class precision and recall results plotted as bar graphs, with dashed lines indicating values expected from either ideal (green) or random (red) sorting.

**Supplementary Figure 2.**
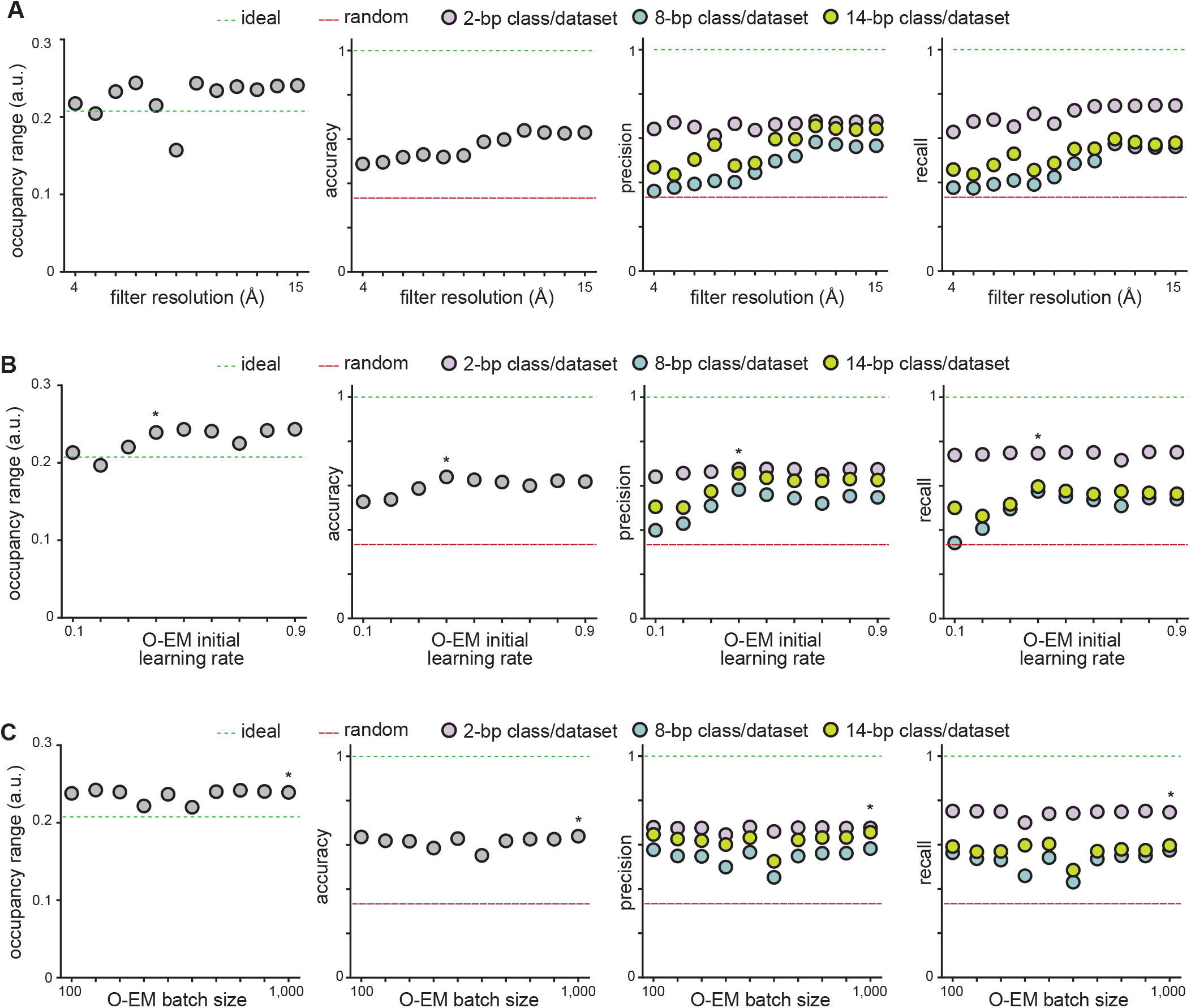
3D classification hyperparameter titrations. **(A-C)** Results from focused 3D classification on the 2/8/14-bp mixed particle stack, with titrated values (see Methods) provided for filter resolution (A), online expectation maximization (O-EM) initial learning rate (B), or O-EM batch size (C). In (B) and (C), an asterisk (*) denotes default values. Dashed lines indicate expected performance under random (red) or ideal (green) particle sorting.

**Supplementary Figure 3.**
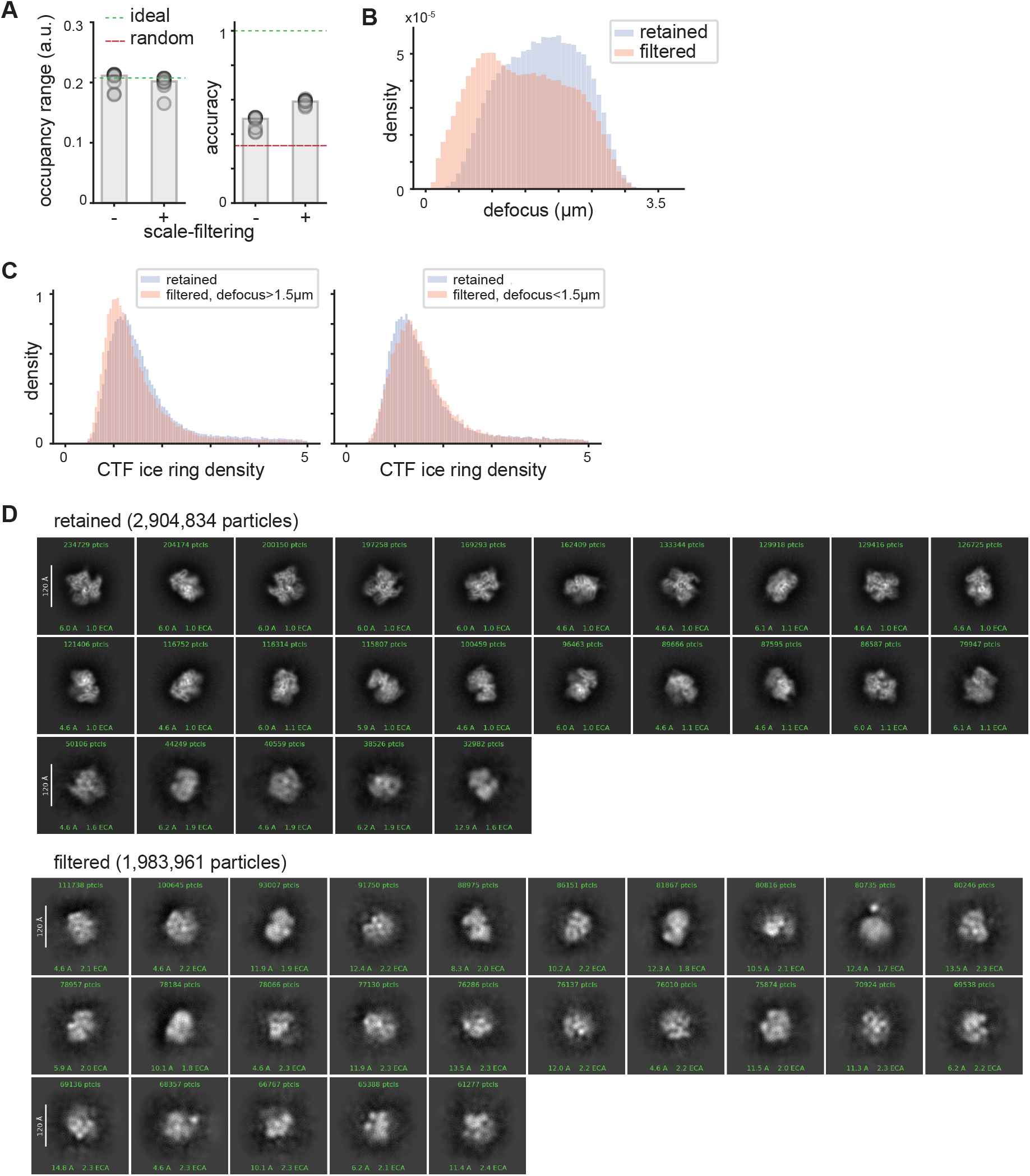
Filtering by per-particle scale factor increases particle sorting accuracy in focused 3D classification. **(A)** Occupancy range of the tDNA extension and particle sorting accuracy assessing performance of focused 3D classification either with or without scale-filtering applied to the particle states, plotted as bar charts (median values) with each dot denoting one of ten replicate classifications. Dashed lines indicate values of each metric expected from either ideal (green) or random (red) sorting. Note that classifications used poses from the original RELION refinement rather than the poses from deep classification used elsewhere in this paper. **(B)** Overlaid histograms depicting defocus distributions (B) or FSC-based ice ring densities estimated by RELION (C) from particles across all datasets that were either retained (blue) or eliminated (peach) during scale-filtering. Distributions for filtered particles are shown conditioned on the particle’s defocus value. **(D)** 2D classes arising from particles that were either retained (top) or eliminated (bottom) during scale-filtering across all 13 datasets spanning the 2-bp to 14-bp tDNA extensions.

**Supplementary Figure 4.**
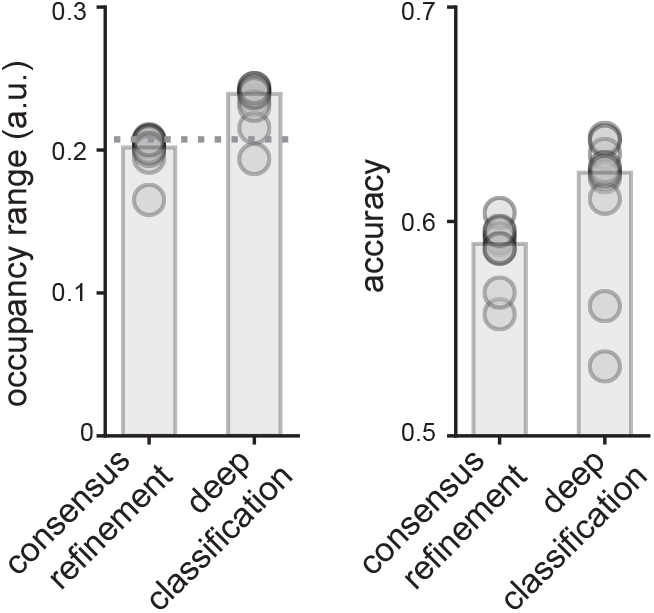
Pose estimation impacts classification accuracy. Occupancy range of the tDNA extension (left) and particle sorting accuracy (right) assessing focused 3D classification using either poses from the original RELION consensus refinement or the poses from deep classification used elsewhere in this paper. Bars represent median values; dots denote each of 10 replicate 3D classifications, and dashed line indicates expected occupancy range based on ideal particle sorting.

**Supplementary Figure 5.**
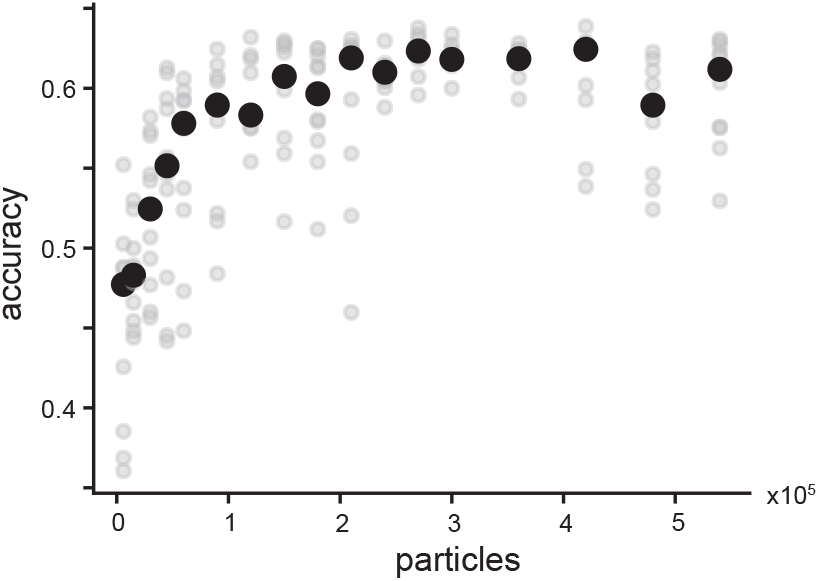
Classification accuracy improves with particle stack size. Scatter plot depicting particle sorting accuracy from focused 3D classification on ten 0.5 replicate particle stacks as a function of the particle stack size. Accuracies for individual replicates are indicated by translucent gray markers; median accuracy value at each particle stack size is indicated by a solid black marker.

**Supplementary Figure 6.**
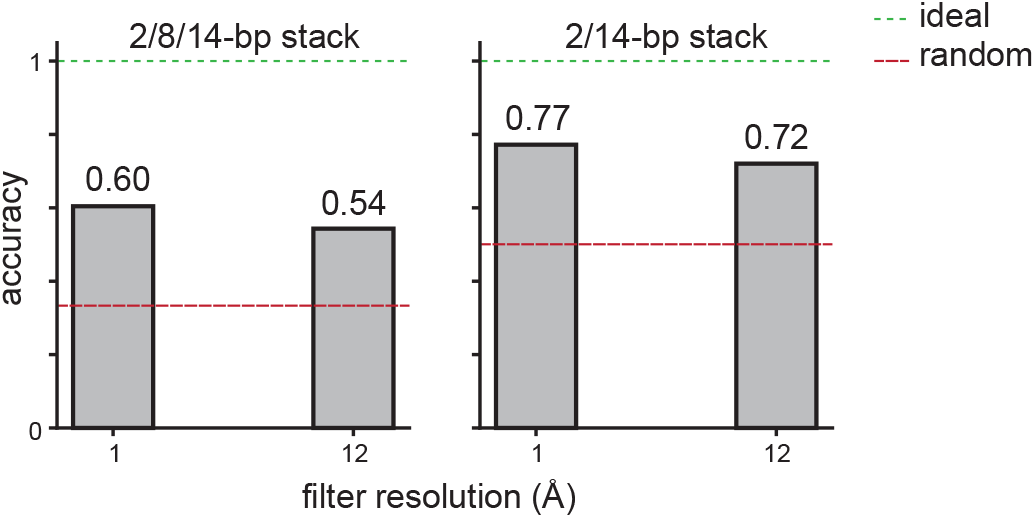
Particle sorting accuracy of single-round classification against correct volumes. Bar charts depicting particle sorting accuracy following a single full expectation-maximization iteration with provided ground-truth volumes (see Methods). Results on different mixed particle stacks and using different filter resolution parameters are shown, with dashed lines indicating performance expected under random (red) and ideal (green) particle sorting.

**Supplementary Figure 7.**
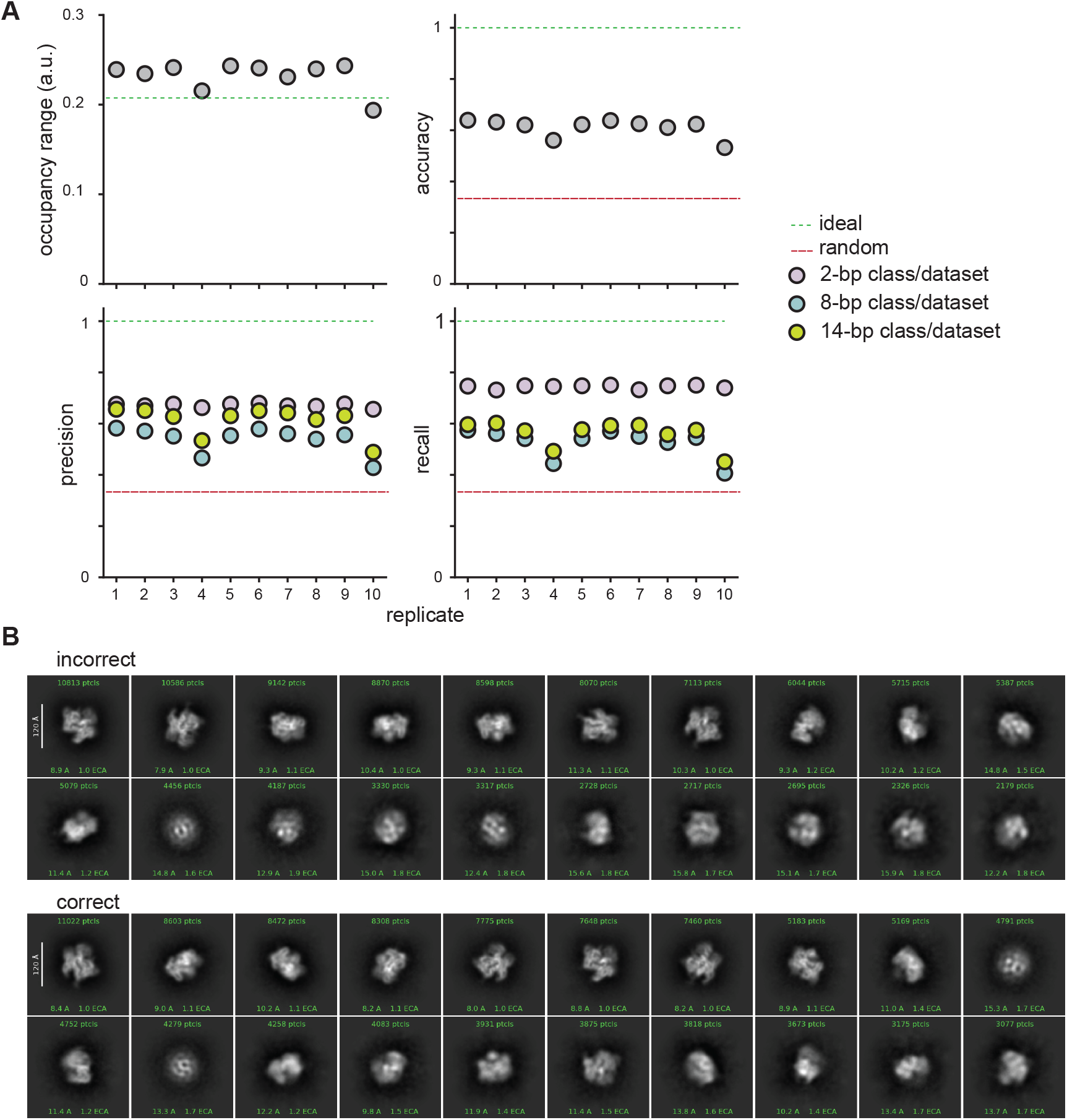
Replicate-to-replicate variability in focused 3D classification on the 2/8/14-bp mixed particle stack. **(A)** Scatter plot of the tDNA extension occupancy range, accuracy, precision, and recall classification metrics across 10 random seed replicates. Dashed lines indicate performance expected under random (red) and ideal (green) particle sorting **(B)** 2D class averages of particles from the 2/8/14-bp mixed particle stack that were always incorrectly (top) or always correctly (bottom) classified by focused 3D classification. The correctly classified particles were randomly subsampled to match the number of particles in the incorrectly classified stack.

**Supplementary Figure 8.**
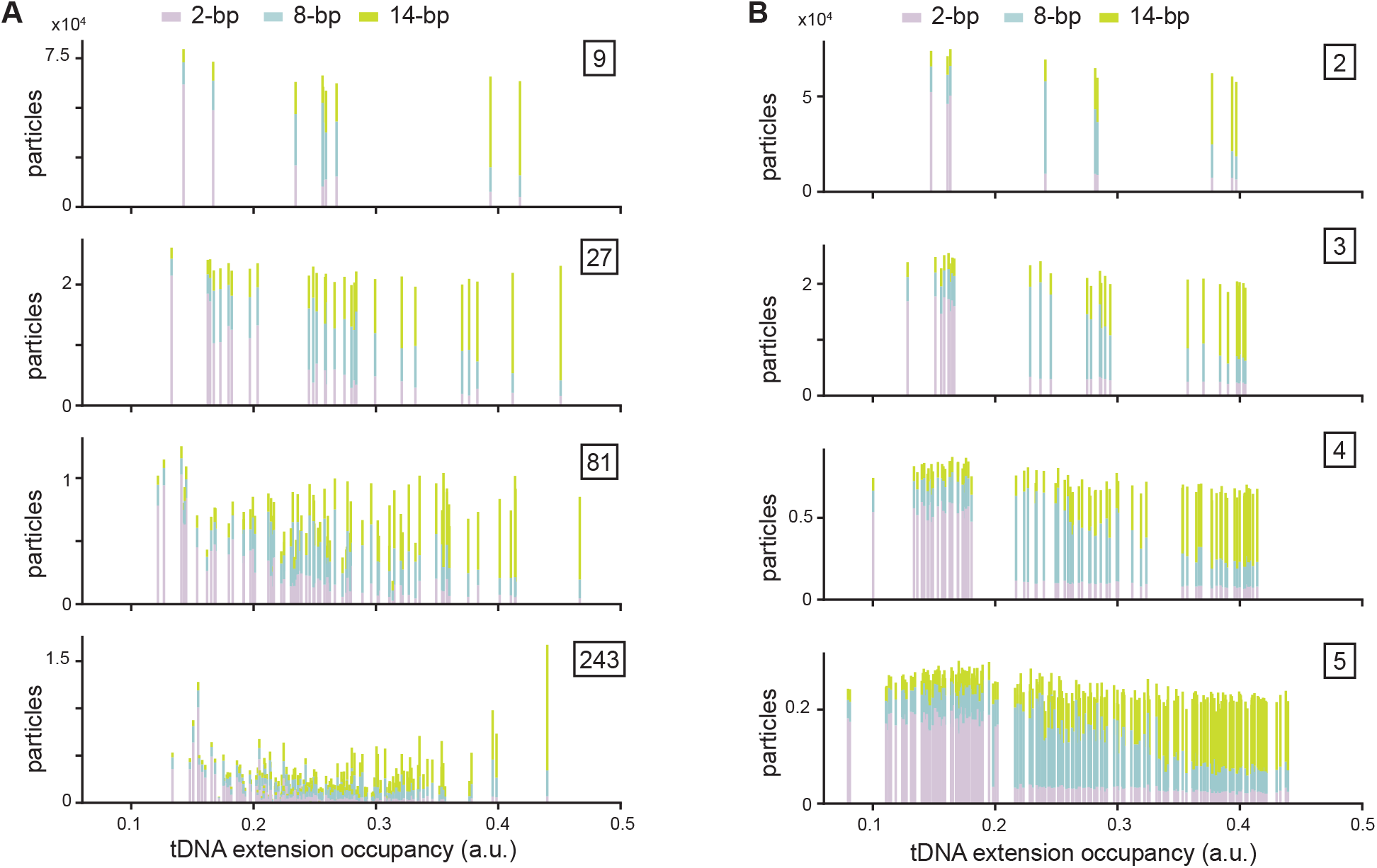
Wide and deep classification with PCA initialization mode. **(A-B)** Stacked bar charts showing the distribution of particles assigned to each class as a function of tDNA extension occupancy when classified using PCA initialization mode (see Methods). Bars are colored according to the ground-truth tDNA extension label (dataset-of-origin). Panel (A) shows results from the wide classification approach, with boxed numbers indicating the total number of classes, while (B) shows results from successive layers of deep classification, with classification layer annotated in box. Each stacked bar represents one class, with mixed-color bars highlighting inaccurate particle classification.

**Supplementary Figure 9.**
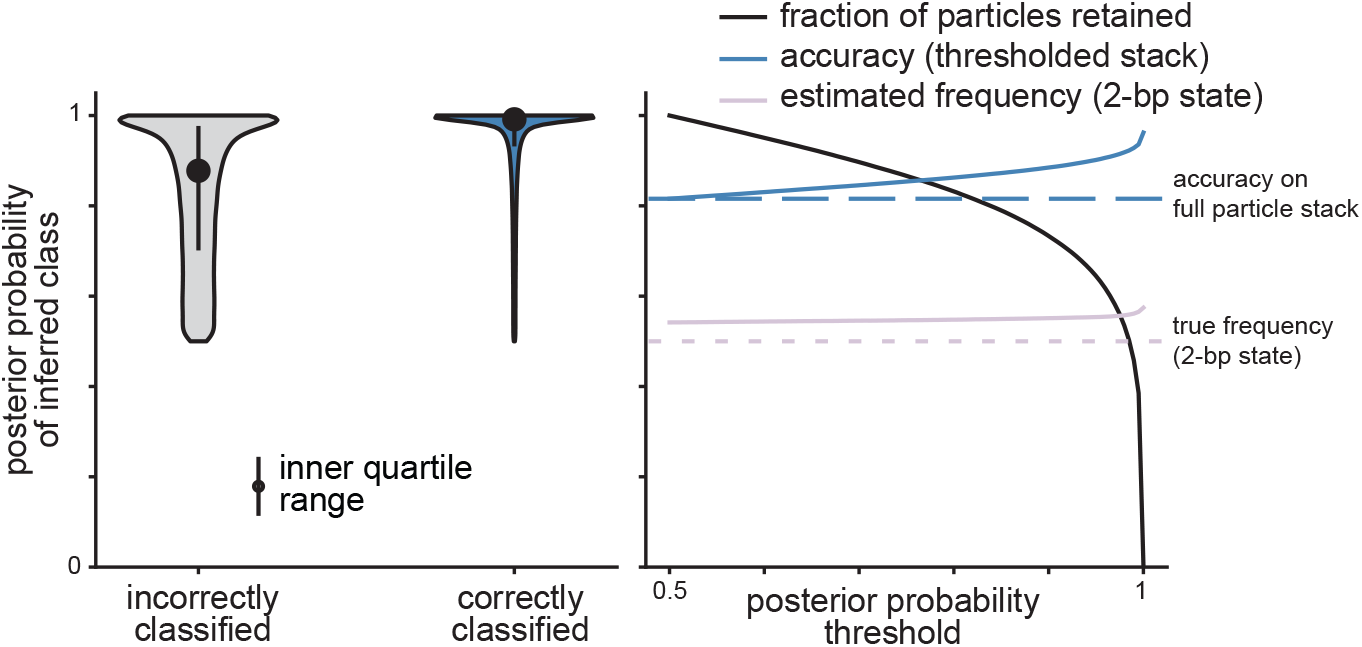
Incorrectly classified particles are sorted less confidently in the 2/14-bp mixed particle stack. Violin plot depicting distribution of per-particle posterior probabilities estimated by focused 3D classification on the 2/14-bp mixed particle stack, conditioned on whether particles were correctly or incorrectly classified (left). Accuracy and estimated frequency of the 2-bp state are plotted as a function of the lower bound imposed on the posterior probability for accepted particle assignments (right). True 2-bp frequency and accuracy on the full particle stack are noted with dashed lines.

**Supplementary Figure 10.**
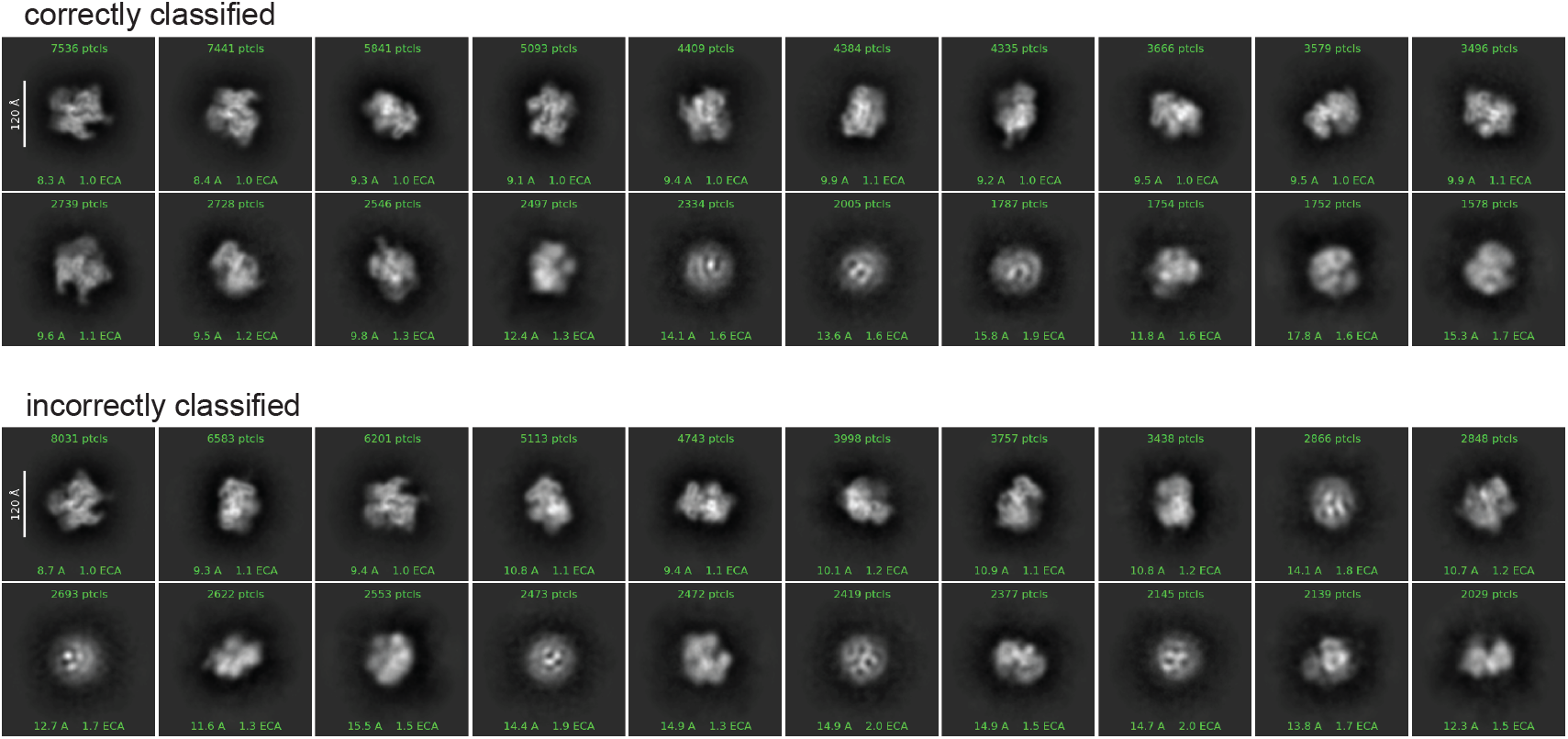
Incorrectly classified particles contain *bona fide* dCas9 particles. 2D class averages from particles in the 2/14-bp mixed particle stack that were always correctly classified (top) or always incorrectly classified (bottom). Correctly classified particles were randomly subsampled to match the number of particles in the incorrectly classified stack.

**Supplementary Figure 11.**
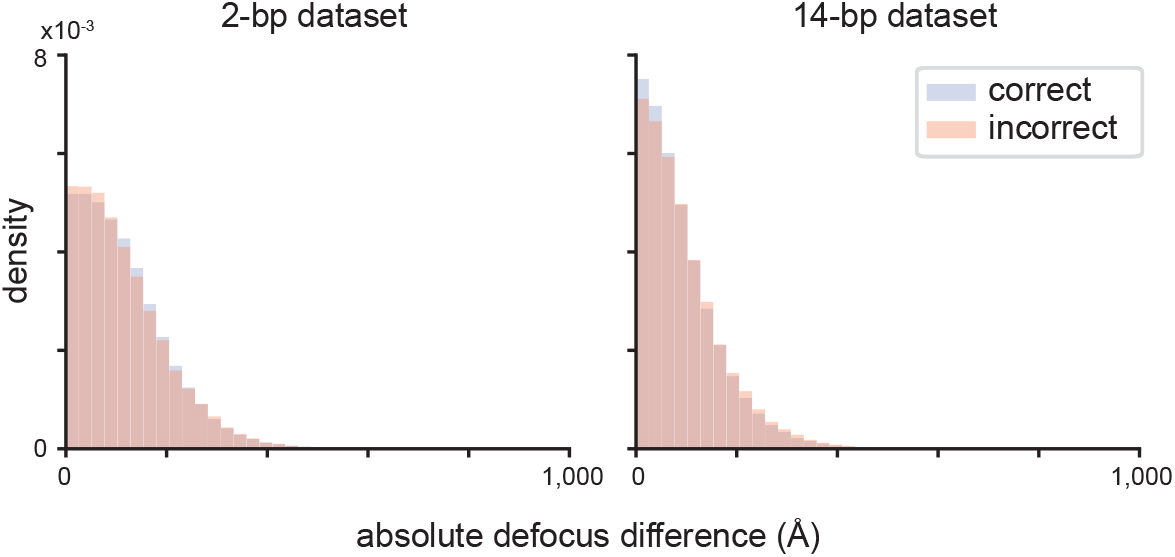
Differences between initial micrograph-level and final per-particle defocus estimates do not correlate with classification accuracy. Overlaid histograms showing the distribution of the absolute difference between the initial micrograph-level defocus estimate and the final per-particle defocus estimate after CTF refinement. Particles are grouped by whether they were always correctly classified (blue) or always incorrectly classified (peach). Distributions are shown for the 2-bp dataset (left) and 14-bp dataset (right).

**Supplementary Figure 12.**
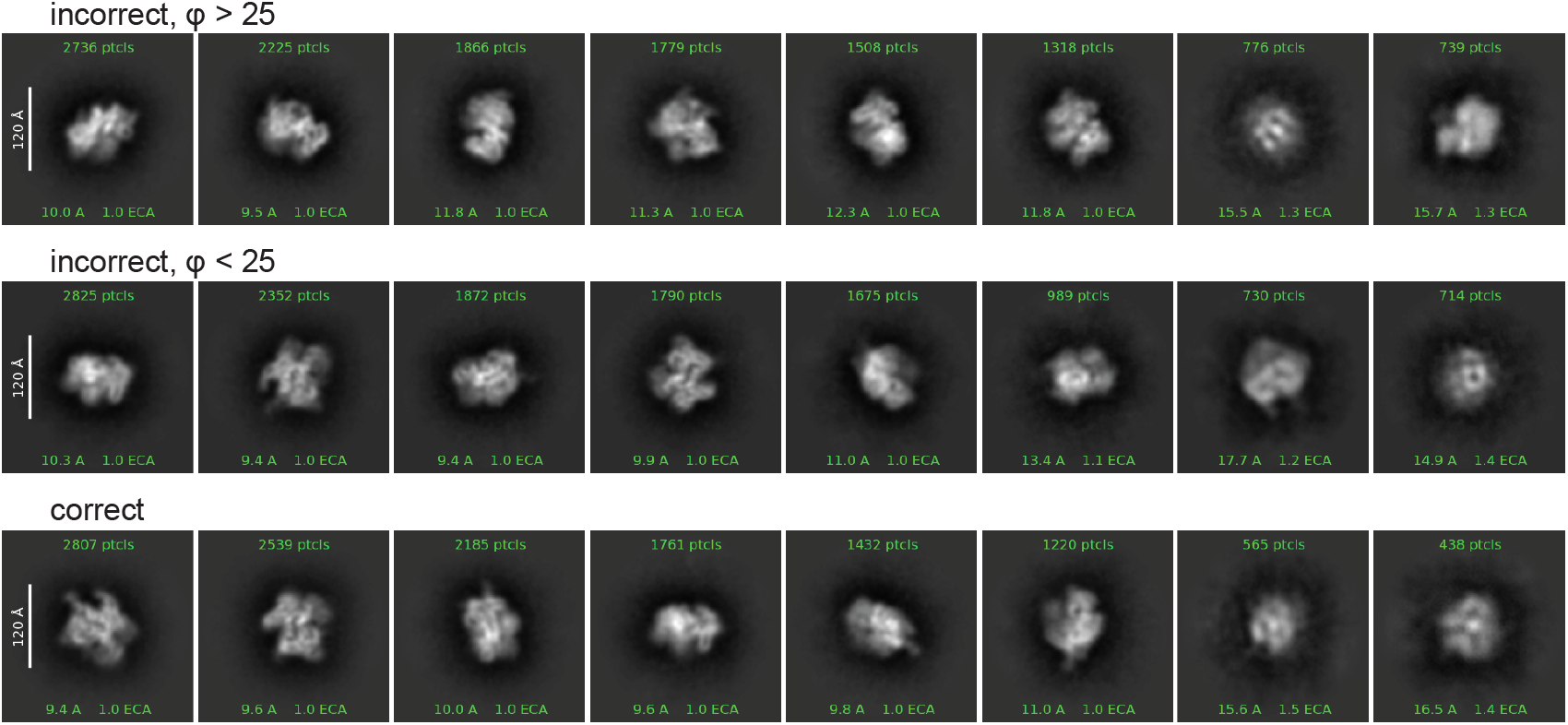
2D class averages of particles from different estimated viewing angles. 2D class averages of particles that were always misclassified in 2/14-bp stack, binned by φ value (top, middle rows) as noted in group label, and those that were always correctly classified (bottom row). Stacks containing misclassified particles with φ < 25 and correctly classified particles were randomly subsampled to match the number of particles in the misclassified φ > 25 stack.

**Supplementary Figure 13.**
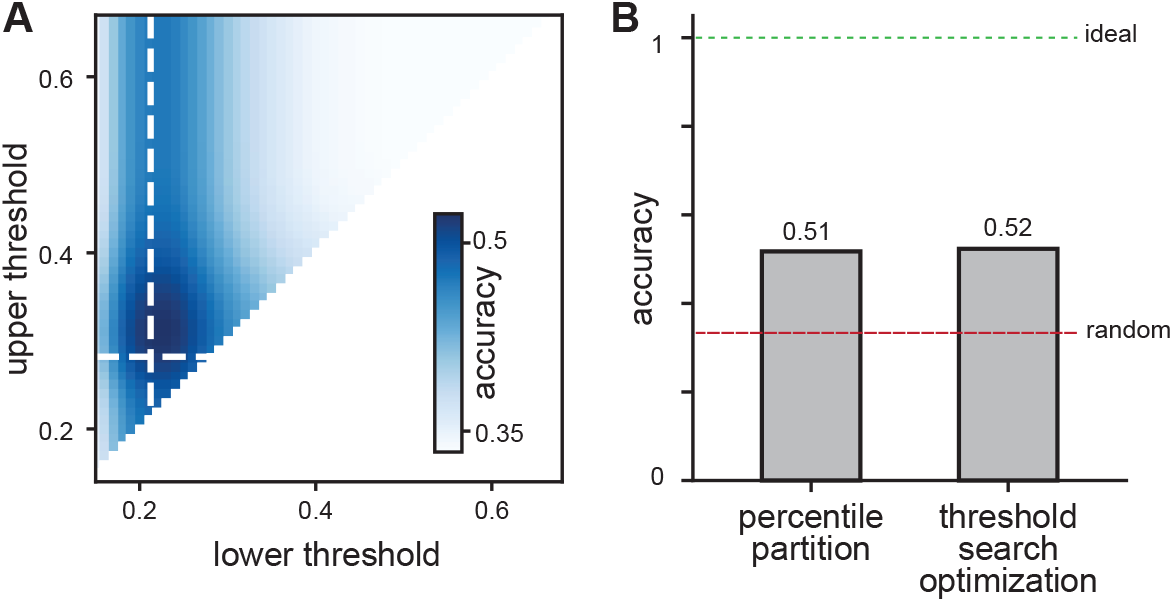
CryoDRGN classification accuracy is robust to chosen occupancy threshold values. **(A)** Heatmap depicting particle sorting accuracy as a function of titrated values for lower (separating 2 bp and 8 bp classes) and upper (separating 8 bp and 14 bp classes) occupancy thresholds (see Methods). Dashed white lines indicate effective threshold values from percentile partitioning approach used elsewhere in this manuscript. **(B)** Bar chart depicting observed accuracy from percentile partitioning approach and maximum achievable accuracy based on systematic threshold search. Dashed lines indicate performance expected under random (red) and ideal (green)

**Supplementary Figure 14.**
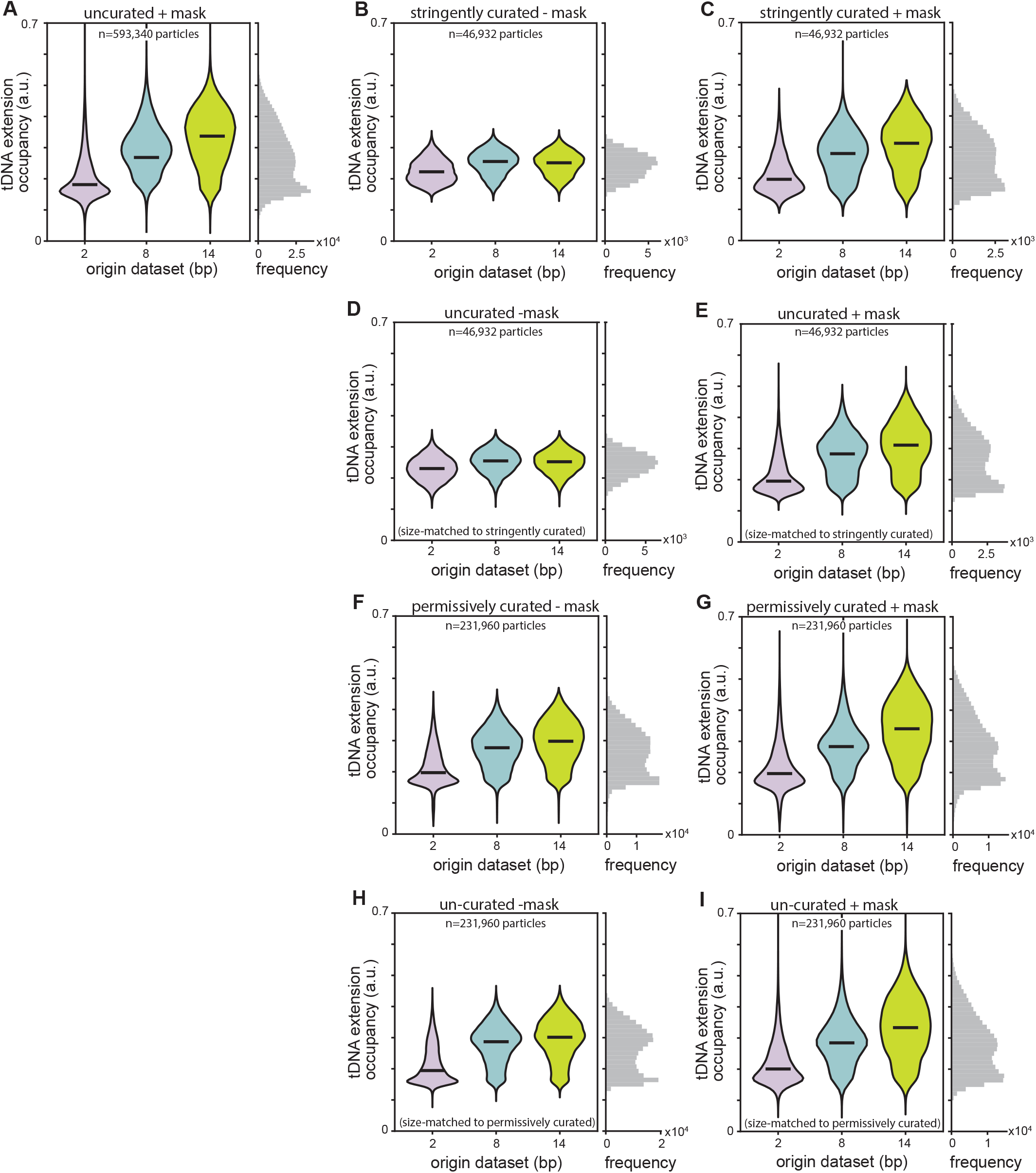
Impact of particle curation and masking on cryoDRGN reconstructions of tDNA extensions. **(A-I)** Violin plots showing distribution of tDNA extension occupancies measured from per-particle volumes generated by cryoDRGN models trained on: the uncurated 2/8/14-bp mixed particle stack (n=593,340 particles), with a mask (A); the stringently curated subset (n=46,932 particles), without a mask (B) and with mask (C); a random subset of the uncurated particle stack (n=46,932 particles), without a mask (D) and with a mask (E); the permissively curated subset (n=231,960 particles), without mask (F) and with mask (G); a random subsample of the uncurated particle stack (n=231,960 particles), without mask (H) and with mask (I).

**Supplementary Figure 15.**
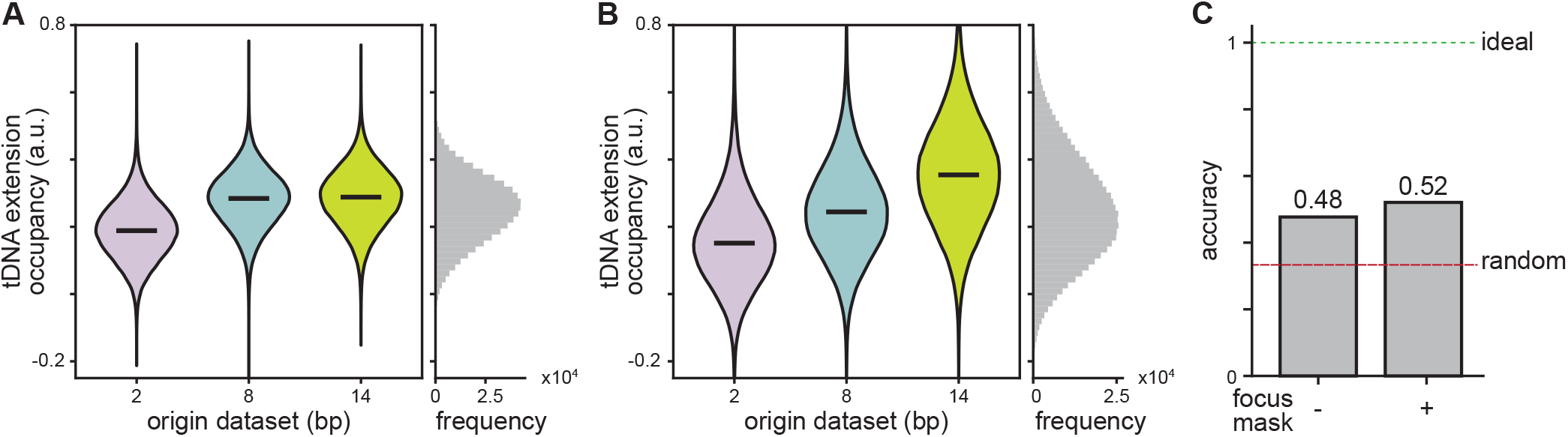
Focused masking modestly improves reconstruction fidelity and particle sorting accuracy in 3D Variability Analysis. **(A-B)** Violin plots showing distribution of tDNA extension occupancies measured from per-particle volumes generated by 3D Variability Analysis of the 2/8/14-bp mixed particle stack with a mask covering (A) the entire volume, or (B) the tDNA extension. **(C)** Bar chart depicting particle sorting accuracy of 3D Variability Analysis with or without a focused mask on the tDNA extension. Dashed lines indicate performance expected under random (red) and ideal (green) particle sorting.

**Supplementary Figure 16.**
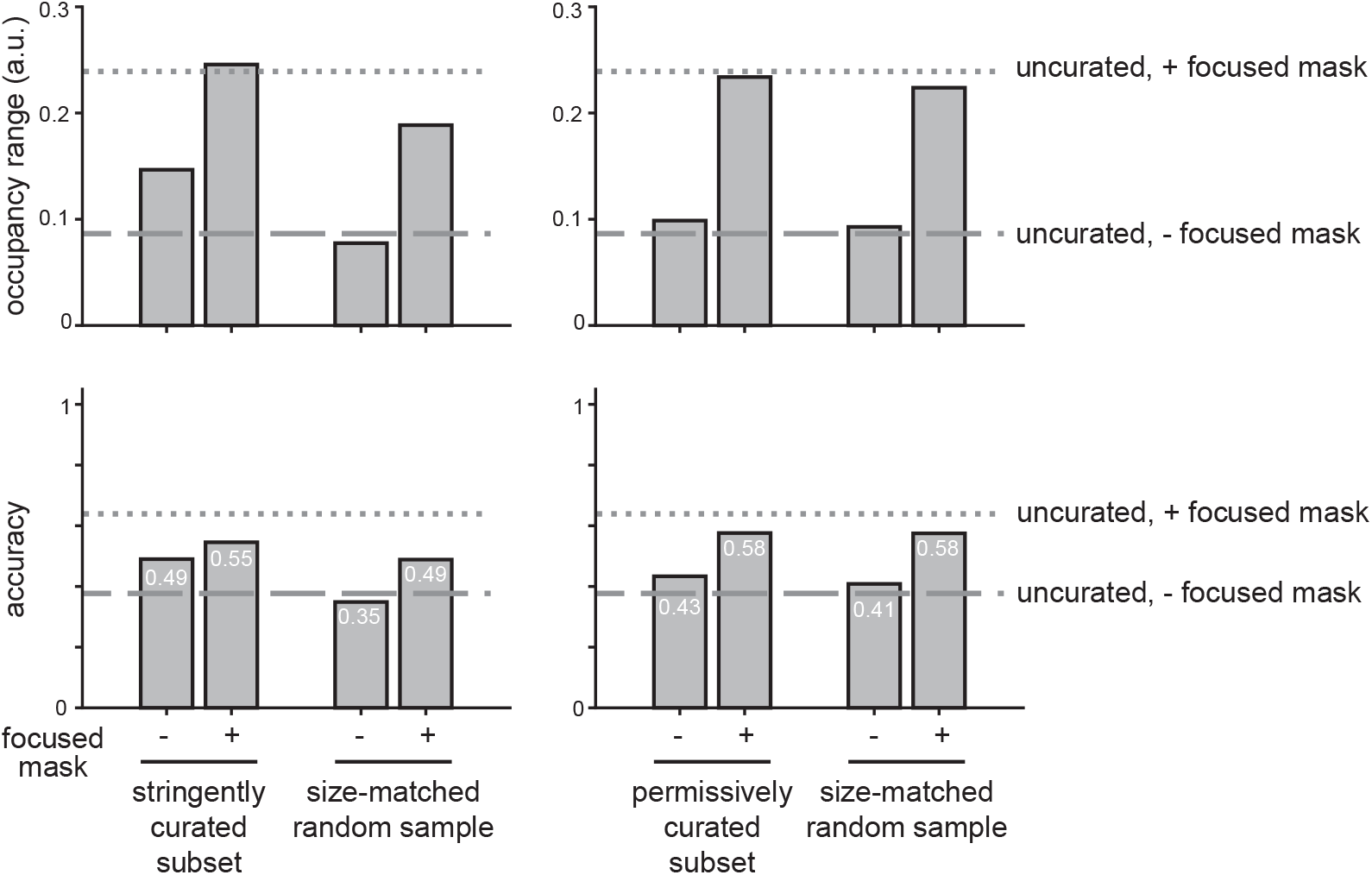
Depleting non-encoded heterogeneity has limited impact on particle sorting accuracy. tDNA extension occupancy range (top) and particle sorting accuracy (bottom) performance metrics plotted as bar charts from focused classification on curated subsets and size-matched randomly sampled controls, with and without a focused mask on the DNA extension. Dashed lines indicate performance on the full, uncurated 2/8/14-bp mixed particle stack with (dotted) or without (dashed) a mask focused on the tDNA extension.

## REFERENCES

Amann SJ, Keihsler D, Bodrug T, Brown NG, Haselbach D. 2023. Frozen in time: analyzing molecular dynamics with time-resolved cryo-EM. Structure 31: 4–19.

Axelrod JJ, Zhang JT, Petrov PN, Glaeser RM, Müller H. 2024. Modern approaches to improving phase contrast electron microscopy. Current Opinion in Structural Biology 86: 102805.

Callaway E. 2015. The revolution will not be crystallized: a new method sweeps through structural biology. Nature 525: 172–174.

Chen JS, Dagdas YS, Kleinstiver BP, Welch MM, Sousa AA, Harrington LB, Sternberg SH, Joung JK, Yildiz A, Doudna JA. 2017. Enhanced proofreading governs CRISPR–Cas9 targeting accuracy. Nature 550: 407–410.

Chen M, Ludtke SJ. 2021. Deep learning-based mixed-dimensional Gaussian mixture model for characterizing variability in cryo-EM. Nature Methods 2021 18:8 18: 930–936.

Cheng Y. 2018. Single particle cryo-EM – how did it get here and where will it go. Science 361: 876–880.

Danev R, Baumeister W. 2016. Cryo-EM single particle analysis with the Volta phase plate ed. S.H. Scheres. eLife 5: e13046.

Danev R, Buijsse B, Khoshouei M, Plitzko JM, Baumeister W. 2014. Volta potential phase plate for in-focus phase contrast transmission electron microscopy. Proceedings of the National Academy of Sciences of the United States of America 111: 15635–15640.

Davis JH, Tan YZ, Carragher B, Potter CS, Lyumkis D, Williamson JR. 2016. Modular Assembly of the Bacterial Large Ribosomal Subunit. Cell 167.

Dingeldein L, Silva-Sánchez D, Evans L, D’Imprima E, Grigorieff N, Covino R, Cossio P. 2025. Amortized template matching of molecular conformations from cryoelectron microscopy images using simulation-based inference. Proceedings of the National Academy of Sciences 122: e2420158122.

Evans L, Dingeldein L, Covino R, Gilles MA, Thiede E, Cossio P. 2026. Counting particles in cryo-electron microscopy may result in incorrect population estimates. Commun Biol 9: 421.

Frank J. 2018. New Opportunities Created by Single-Particle Cryo-EM: The Mapping of Conformational Space. Biochemistry 57: 888–888.

Ghanbarpour A, Cohen SE, Fei X, Kinman LF, Bell TA, Zhang JJ, Baker TA, Davis JH, Sauer RT. 2023. A closed translocation channel in the substrate-free AAA+ ClpXP protease diminishes rogue degradation. Nature Communications 2023 14:1 14: 1–10.

Gilles MA, Singer A. 2025. Cryo-EM heterogeneity analysis using regularized covariance estimation and kernel regression. Proceedings of the National Academy of Sciences 122: e2419140122.

Grassetti AV, Kinman LF, Davis JH. 2026. Engineered Cas9 complexes establish an experimentally grounded benchmark for heterogeneous cryoEM reconstruction methods. bioRxiv biorxiv.org/content/10.64898/2026.05.04.721978v1.

Haselbach D, Schrader J, Lambrecht F, Henneberg F, Chari A, Stark H. 2017. Long-range allosteric regulation of the human 26S proteasome by 20S proteasome-targeting cancer drugs. Nature communications 8.

Jeon M, Raghu R, Astore M, Woollard G, Feathers R, Kaz A, Hanson SM, Cossio P, Zhong ED. 2024. CryoBench: Diverse and challenging datasets for the heterogeneity problem in cryo-EM. arXiv arxiv.org/abs/2408.05526.

Kinman LF, Carreira MV, Powell BM, Davis JH. 2025. Automated model-free analysis of cryo-EM volume ensembles with SIREn. Structure 33: 974-987. e4.

Kinman LF, Powell BM, Zhong ED, Berger B, Davis JH. 2023. Uncovering structural ensembles from single-particle cryo-EM data using cryoDRGN. Nature Protocols 18: 319–339.

Lederman RR, Singer A. 2017. Continuously heterogeneous hyper-objects in cryo-EM and 3-D movies of many temporal dimensions. arXiv arxiv.org/abs/1704.02899.

Leschziner A. 2010. The Orthogonal Tilt Reconstruction Method. In Methods in Enzymology, Vol. 482 of, pp. 237–262.

Lyumkis D, Brilot AF, Theobald DL, Grigorieff N. 2013. Likelihood-based classification of cryo-EM images using FREALIGN. Journal of Structural Biology 183: 377–388.

May MB, Lopez-Perez GS, Davis JH. 2026. Capturing ribosomal structures in cellular extracts with cryoPRISM: A purification-free cryoEM approach reveals novel structural states. Proc Natl Acad Sci U S A 123: e2521210123.

Meng EC, Goddard TD, Pettersen EF, Couch GS, Pearson ZJ, Morris JH, Ferrin TE. 2023. UCSF ChimeraX: Tools for structure building and analysis. Protein Science 32: e4792.

Penczek PA, Kimmel M, Spahn CMT. 2011. Identifying Conformational States of Macromolecules by Eigen-Analysis of Resampled Cryo-EM Images. Structure 19: 1582–1590.

Plaschka C, Lin P-C, Nagai K. 2017. Structure of a pre-catalytic spliceosome. Nature 546: 617–621.

Powell BM, Davis JH. 2024. Learning structural heterogeneity from cryo-electron sub-tomograms with tomoDRGN. Nature Methods 2024 1–12.

Punjani A, Fleet DJ. 2021. 3D variability analysis: Resolving continuous flexibility and discrete heterogeneity from single particle cryo-EM. Journal of Structural Biology 213: 107702.

Punjani A, Fleet DJ. 2023. 3DFlex: determining structure and motion of flexible proteins from cryo-EM. Nature Methods 2023 20:6 20: 860–870.

Punjani A, Rubinstein JL, Fleet DJ, Brubaker MA. 2017. cryoSPARC: algorithms for rapid unsupervised cryo-EM structure determination. Nature Methods 2017 14:3 14: 290–296.

Rabuck-Gibbons JN, Lyumkis D, Williamson JR. 2022. Quantitative Mining of Compositional Heterogeneity in Cryo-EM Datasets of Ribosome Assembly Intermediates. Structure (London, England: 1993) 30: 498.

Remis J, Petrov PN, Zhang JT, Axelrod JJ, Cheng H, Sandhaus S, Mueller H, Glaeser RM. 2024. Cryo-EM phase-plate images reveal unexpected levels of apparent specimen damage. Journal of Structural Biology 216: 108150.

Rosenthal PB, Henderson R. 2003. Optimal determination of particle orientation, absolute hand, and contrast loss in single-particle electron cryomicroscopy. J Mol Biol 333: 721–745.

Scheres SHW. 2012. RELION: Implementation of a Bayesian approach to cryo-EM structure determination. Journal of Structural Biology 180: 519.

Scheres SHW, Valle M, Nuñez R, Sorzano COS, Marabini R, Herman GT, Carazo JM. 2005. Maximum-likelihood Multi-reference Refinement for Electron Microscopy Images. Journal of Molecular Biology 348: 139–149.

Schwab J, Kimanius D, Burt A, Dendooven T, Scheres SHW. 2024. DynaMight: estimating molecular motions with improved reconstruction from cryo-EM images. Nat Methods 21: 1855–1862.

Schwartz O, Axelrod JJ, Campbell SL, Turnbaugh C, Glaeser RM, Müller H. 2019. Laser phase plate for transmission electron microscopy. Nat Methods 16: 1016–1020.

Sigworth FJ. 2016. Principles of cryo-EM single-particle image processing. Microscopy 65: 57.

Sun J, Kinman LF, Jahagirdar D, Ortega J, Davis JH. 2023. KsgA facilitates ribosomal small subunit maturation by proofreading a key structural lesion. Nature Structural & Molecular Biology 2023 1–13.

Tagare HD, Kucukelbir A, Sigworth FJ, Wang H, Rao M. 2015. Directly reconstructing principal components of heterogeneous particles from cryo-EM images. Journal of Structural Biology 191: 245–262.

Tang WS, Silva-Sánchez D, Giraldo-Barreto J, Carpenter B, Hanson SM, Barnett AH, Thiede EH, Cossio P. 2023. Ensemble Reweighting Using Cryo-EM Particle Images. J Phys Chem B 127: 5410–5421.

Toader B, Sigworth FJ, Lederman RR. 2023. Methods for Cryo-EM Single Particle Reconstruction of Macromolecules Having Continuous Heterogeneity. J Mol Biol 435: 168020.

Zhong ED, Bepler T, Berger B, Davis JH. 2021. CryoDRGN: reconstruction of heterogeneous cryo-EM structures using neural networks. Nature Methods 18.

